# Polymerization of Tetraspanin 7 into Helical Transmembrane Skeletons for Tubular Membrane Stabilization

**DOI:** 10.1101/2024.06.27.600806

**Authors:** Dongju Wang, Xia Jia, Raviv Dharan, Jiahao Ren, Yi Zheng, Xiaopeng Li, Mingtao Huang, Kui Xu, Qi Zhang, Takami Sho, Siyuan Liu, Fan Yang, Qiangfeng Cliff Zhang, Raya Sorkin, Nan Liu, Hong-Wei Wang, Li Yu

## Abstract

Tubular cell membrane protrusions, such as filopodia^1^, dendrites^2^, tunneling nanotubes (TNTs)^3^ and retraction fibers^4^, are prevalent on the cell surface. They play crucial roles in various physiological processes and disease states, including angiogenesis, wound healing, and cancer metastasis^5–7^. The shaping and stabilization of these protrusions are crucial for their function, and the molecular machineries involved have long been an active area of investigation. Here we employed an integrative approach combining live-cell imaging, *in vitro* reconstitution, and *in situ* cryo-EM analysis to reveal that the transmembrane protein tetraspanin 7 (TSPAN7) senses membrane curvature and polymerizes into a helical configuration on highly curved tubular membranes. These spirals act as a “transmembrane skeleton”, effectively maintaining the structural integrity of membrane protrusions under mechanical stress. Our findings reveal a previously unreported assembly strategy for transmembrane proteins and a mechanism for maintaining tubular membrane protrusions.

## Main

Tubes are a fundamental structural unit of cellular organization. Tubular cell membrane protrusions of various scales connect spatially distant parts of the cell body, enabling communication and coordination, which significantly enhance tissue organization complexity^5^. Neurons feature specialized tubular membrane protrusions, including axons, dendrites, and spines^2^. Other tubular membrane protrusions, such as cilia^8^, filopodia, tunneling nanotubes, cytonemes^9^, and retraction fibers, are prevalent across diverse cell types.

The shaping and stabilization of tubular membrane protrusions are essential and complicated processes. They involve the initiation of high curvature from a flat membrane, which requires overcoming a substantial energy barrier^10,11^. While the flexibility of tubular membrane protrusions allows for dynamic functions, it comes at the expense of fragility. The cell surface encounters a variety of mechanical stresses: fluid flow generates shear force, while adhesion between cells or between a cell and the extracellular matrix (ECM) creates traction forces^12^. These forces can induce bending or narrowing of tubular membranes, potentially leading to their rupture. This underscores the importance for structural stability and the need for mechanisms to preserve it^13^. The cytoskeleton plays a central role in forming and maintaining tubular membrane protrusions, such as microtubule-based structures like cilia and axons^14,15^, as well as filamentous actin (F-actin)-supported filopodia^1^. Additionally, proteins like spectrins link the membrane to actin, further contributing to the stability of these structures^16^. However, it remains an open question whether other mechanisms exist to support tubular membrane protrusions.

Recent studies have underscored the involvement of tetraspanins (TSPANs), a family of four-pass transmembrane proteins, in the formation of tubular protrusions on the plasma membrane^17–20^. Specifically, CD9/TSPAN29 and TSPAN4 were identified as sensors for positive membrane curvature and contribute to the remodeling process of tubular membranes^21^. Tetraspanins are best known as organizers on membranes, and two distinct organizational strategies have been identified. Tetraspanins can assemble into tetraspanin-enriched microdomains (TEMs) by recruiting various protein partners and lipids^22^. Additionally, tetraspanin oligomers are capable of self-organizing into highly ordered supramolecular assemblies. For instance, TSPAN22/Peripherin 2 and TSPAN23/ROSP1 form linear polymers and shape the rim of the retinal photoreceptor outer segment disks^23^, and TSPAN21/UPK1A and TSPAN20/UPK1B form hexagonal oligomers that further assemble into 2D crystal-like arrays on urothelial plaques^24,25^.

TSPAN7 is a member of the tetraspanin family and has been implicated in multiple diseases such as intellectual disability^26–28^, viral infection^29^, diabetes^30^ and cancer development^31,32^. Overexpression of TSPAN7 promotes the formation of filopodia and dendritic spines in hippocampal neurons, and knockdown of TSPAN7 enhances spine motility and turnover^33^. Moreover, TSPAN7 expression was induced during the maturation of monocyte-derived dendritic cells (MDDCs). Knockdown of TSPAN7 impairs dendrite formation in MDDCs and blocks dendrite-mediated transfer of HIV-1 to T cells^29^. These functional studies of TSPAN7 highlight its involvement in shaping and stabilizing tubular membranes. However, the underlying molecular mechanism remains largely elusive.

In this study, we reveal that TSPAN7 molecules sense high membrane curvature and self-assemble into a helical configuration on tubular membranes. In the helical assembly, TSPAN7 protomers organize through head-to-head and back-to-back interactions to form a spiral protofilament, and multiple protofilaments wind together to stabilize the tubular membrane protrusions under mechanical stress by resisting the stretching-induced narrowing and breakage. The TSPAN7 helical arrangement is inversely correlated with the presence of actin fibers in the tubular membrane protrusion, suggesting a complementary mechanism for membrane tubule stabilization. Our findings lead us to introduce the concept of the “transmembrane skeleton”, a structural arrangement of membrane proteins that shapes and stabilizes tubular protrusions via a previously unrevealed mechanism.

### TSPAN7 senses high curvature

In order to assess the subcellular localization and dynamics of TSPAN7 in cells, we expressed GFP-tagged rat TSPAN7 in normal rat kidney (NRK) cells. Compared to control GFP vector, the expression of TSPAN7-GFP significantly enhanced the formation of retraction fibers and TNTs – two typical types of tubular membrane protrusions that are generated via cell-ECM and cell-cell interactions, respectively (Fig. 1a, b, c). Compared to TNTs, retraction fibers are much easier to observe under a fluorescence microscope. Therefore, we chose retraction fibers as the model system for the following studies. We constructed cell lines with different expression levels of TSPAN7-GFP and observed retraction fiber formation in these cells. The results showed that TSPAN7-GFP promotes the formation of retraction fibers in a dose-dependent manner (Extended Data Fig. 1).

**Fig. 1.**
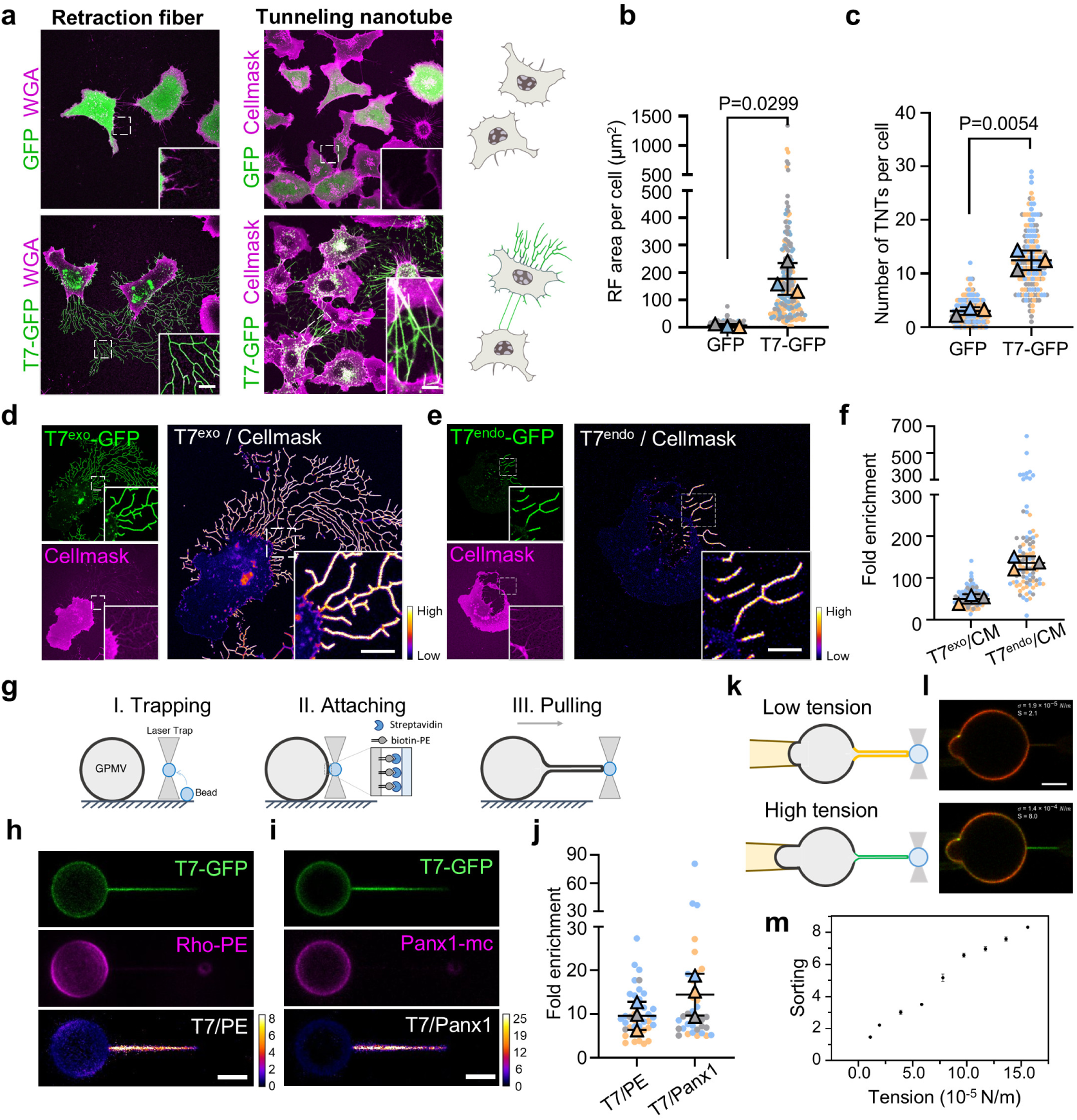
TSPAN7 senses high curvature and promotes the formation of tubular protrusions. **a,** Representative confocal images of retraction fibers (left panel) and tunneling nanotubes (right panel) generated by NRK cells expressing GFP or TSPAN7-GFP (T7-GFP). Cells were stained with tetramethylrhodamine-conjugated WGA (WGA) or CellMask Deep Red (Cellmask). **b,** Quantification of the area of retraction fibers per cell in **a**. N = 179 and 186 cells were analyzed for GFP and T7-GFP, respectively. See Methods section for details on statistical analysis. **c,** Quantification of the number of TNTs per cell in **a**. N = 159 and 155 cells were analyzed for GFP and T7-GFP respectively. **d,** Representative confocal images showing the distribution of overexpressed T7-GFP (T7^exo^-GFP) and Cellmask in NRK cells. The ratio of T7^exo^-GFP/Cellmask is shown as a heatmap. **e,** Representative confocal images showing the distribution of endogenous TSPAN7-GFP (T7^endo^-GFP) and Cellmask in GFP^+^ cells isolated from the lymph nodes of TSPAN7-GFP knock-in mice. The ratio of T7 ^endo^-GFP/Cellmask is shown as a heatmap. **f,** Statistical analysis of the results of **d** and **e**. The fold enrichment 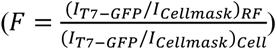 was calculated. N = 125 and 84 cells were analyzed for **d** and **e**, respectively. **g,** Schematic illustration of the GMPV pulling assay. **h, i,** Representative confocal images of GPMVs with a pulled membrane tether. GPMVs were produced by NRK cells expressing T7-GFP and stained with rhodamine-PE (Rho-PE) (**h**) or by NRK cells expressing T7-GFP and PANX1-mcherry (Panx1-mc) (**i**). **j,** Statistical analysis of the results of **h** and **i**. The fold enrichment 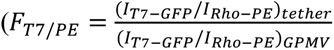 and 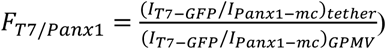 of T7-GFP on the membrane tether versus the GPMV was calculated. N = 38 and 36 GPMVs were analyzed for **h** and **i,** respectively. **k,** Schematic illustration of the aspirated GMPV pulling assay. The membrane tension of the GPMV was gradually increased by raising the aspiration pressure, which increased the curvature of the membrane tether. **l,** Representative confocal images of aspirated GPMV with a membrane tether at low (*σ* = 1.9 × 10^−5^ *N*/*M*, upper panel) and high (*σ* = 1.4 × 10^−5^ *N*/*M* lower panel) membrane tensions. GPMVs were produced from HEK293T cells expressing T7-GFP (green) and stained with DiI-C12 (red). **m,** Representative graph of sorting 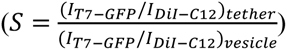 as a function of membrane tension of the GPMV. Error bars are SEM. Scale bars, 5 μm.

We noticed that TSPAN7 is enriched on retraction fibers. In order to perform quantitative analysis of this enrichment, we used cells stably expressing TSPAN7-GFP and we stained the plasma membrane with the dye CellMask Deep Red. When we analyzed their fluorescence intensities on retraction fibers and cell bodies, we found that CellMask showed less partitioning on retraction fibers compared to cell bodies, while TSPAN7 showed nearly 50-fold enrichment on retraction fibers (Fig. 1d, f). We wondered if this enrichment is conserved for endogenous TSPAN7. Due to the lack of commercially available antibodies for TSPAN7, we established a mouse strain where the stop codon of TSPAN7 is replaced by the coding sequence of GFP. The endogenous TSPAN7 then became GFP-tagged (Extended Data Fig. 2a). We sorted GFP^+^ cells from lymph nodes, and identified this population as lymphatic endothelial cells (LECs) (Extended Data Fig. 2b, c, d). We analyzed the distribution of endogenous TSPAN7 and found that the endogenous protein was even more enriched on retraction fibers (136-fold enrichment on average) compared to the exogenously expressed TSPAN7-GFP (Fig. 1e, f).

The preference of TSPAN7 for cylindrical retraction fibers suggests its potential sensitivity to high curvature. We established an *in vitro* system to test this hypothesis (Fig. 1g). The system includes a biotin-PE-labeled giant plasma membrane vesicle (GPMV) with TSPAN7-GFP as a cell-mimetic membrane source and a streptavidin-coated bead as its paired interactor. An optical trap was applied to manipulate the movement of the bead: first approaching and interacting with the GPMV, and then moving in the opposite direction to pull out a cylindrical membrane tether from the GPMV. The radius of the tether was much smaller than that of the GPMV. This results in a continuous membrane system where the curvature of the GPMV is effectively very low, but that of the tether is high.

To examine partitioning of TSPAN7-GFP into the tether, we labeled the membrane with rhodamine-PE (an indicator of bulk lipids) or PANX1-mcherry (a protein with four-transmembrane helices from the non-tetraspanin-related pannexin family) and compared their fluorescence intensities. We found that rhodamine-PE or PANX1-mcherry were almost exclusively localized on GPMVs but nearly undetectable on membrane tethers (Fig. 1h, i). In contrast, TSPAN7-GFP showed a clear preference for localization to the highly curved membrane tethers compared to the almost flat membrane of the GPMVs (Fig. 1h, i, j).

To confirm that TSPAN7 is sensitive to membrane curvature, we integrated micropipette aspiration with optical tweezers. This combination allowed us to control the membrane tension of the GPMV and thus the membrane curvature of the tether^21^. Increasing the aspiration pressure enhances the membrane tension of the GPMV, which can lead to constriction of the tether, i.e. an increase in tether curvature (Fig. 1k). When we pulled the tether at relatively low tension, an enrichment of TSPAN7 in the tether was observed (Fig. 1l). With a gradually increased membrane tension, the tether curvature became higher and the TSPAN7 enrichment in the tether also increased (Fig. 1l, m). These results emphasize the curvature sensitivity of TSPAN7 and suggest that TSPAN7 molecules enrich on retraction fibers due to the high positive membrane curvature of these structures.

### TSPAN7 stabilizes tubular membranes in vitro

Overexpression of TSPAN7-GFP significantly enhanced the formation of retraction fibers and TNTs, with TSPAN7-GFP highly enriched on both types of tubular structures, suggesting that TSPAN7 may promote tubular membrane formation by stabilizing these intrinsically fragile structures. To test this hypothesis, we established a minimized *in vitro* reconstitution system based on synthetic giant unilamellar vesicles (GUVs) and recombinant TSPAN7-GFP protein to avoid cell-derived interferences (Fig. 2a). When contacting a hydrophilic glass surface, a GUV can spontaneously flatten and expand due to strong adhesion, leading to a final rupture at its center. The ruptured edge of the GUV then retracts to the periphery, resulting in the formation of retraction fiber-like membrane tubules. We observed that in protein-free GUVs, this tubular network was unstable and transformed into vesicles within minutes of forming (Fig. 2b, top, and Supplementary video 1). Remarkably, in GUVs embedded with purified TSPAN7-GFP, we observed a rapid ingress of TSPAN7-GFP signal into the tubular membranes (Fig. 2b, bottom, and Supplementary video 2). The tubular network was efficiently stabilized, even after overnight incubation (Fig. 2c). Since TSPAN7-GFP is the sole protein component in this system, these data strongly suggest that TSPAN7 is sufficient for the sensing and stabilization of membrane curvature.

**Fig. 2.**
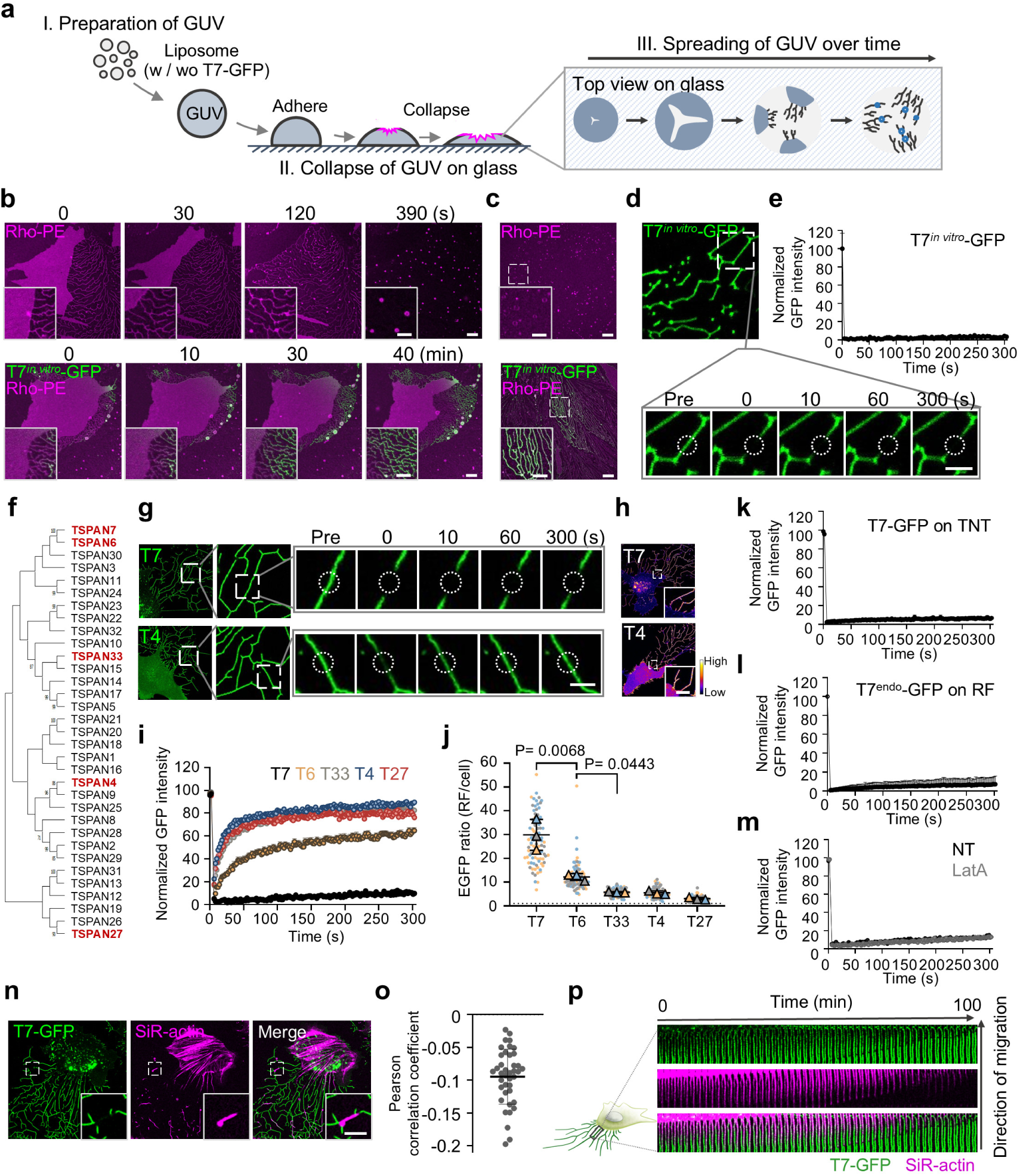
TSPAN7 stabilizes tubular membranes and exhibits immobility on these structures. **a,** Schematic illustration of the GUV-based *in vitro* reconstitution system. **b,** Snapshots from a time-lapse video showing the spreading of a GUV without protein (top panel) or embedded with purified TSPAN7-GFP (T7*^in^ ^vitro^*-GFP, bottom panel). See also **Supplementary video 1** and **Supplementary video 2**. **c,** Representative confocal images showing the morphology of reconstitution systems without protein (top) or with TSPAN7-GFP (bottom) after sitting at room temperature overnight. **d-e,** FRAP assays were performed on *in vitro* reconstituted fibers using TSPAN7-GFP-embedded GUVs. Time-lapse images before and after bleaching are shown in **d**. Dashed circles represent areas of bleaching. The GFP intensities in bleached areas were tracked and plotted in **e**. Data from 9 bleached areas were normalized and plotted as mean + SEM. **f,** Phylogenetic tree of tetraspanin family generated by MEGA11^35^. Five tetraspanins are highlighted in red: TSPAN7; its closest relative TSPAN6; and representative tetraspanins from three different clusters, TSPAN33, TSPAN4 and TSPAN27. **g,** Representative time-lapse images of FRAP assays on the retraction fibers of NRK cells expressing TSPAN7-GFP (T7) or TSPAN4-GFP (T4). The right panels show enlarged areas from the left panels. The dashed circles represent areas of bleaching. **h,** The fluorescence intensities of TSPAN7-GFP (T7) and TSPAN4-GFP (T4) are shown as heatmaps. **i,** The fluorescence intensities in bleached areas in **g** and **Extended Data Fig 3a** were tracked and plotted. For each curve, data from 15 bleached areas were normalized and plotted as mean + SEM. **j,** The ratio of GFP intensity on retraction fibers (RF) versus cell bodies (cell) in **h** and **Extended Data Fig 3b** was analyzed. N = 90 cells were analyzed for each group. **k,** FRAP assays were performed on the TNTs of NRK cells expressing T7-GFP. The GFP intensities in 13 bleached areas were tracked, then normalized and plotted as mean + SEM. See also **Extended Data Fig. 4a**. **l,** FRAP assays were performed on the retraction fibers of GFP^+^ cells isolated from the lymph nodes of TSPAN7-GFP knock-in mice. The GFP intensities in 48 bleached areas were tracked, then normalized and plotted as mean + SEM. See also **Extended Data Fig. 4b**. **m,** FRAP assays were performed on cells pretreated with 2 μM LatA or control solvent (ethanol, NT). For each treatment (LatA or NT), the GFP intensities in 15 bleached areas were tracked, then normalized and plotted as mean + SEM. See also **Extended Data Fig. 4c**. **n,** Representative confocal images of NRK cells stably expressing T7-GFP and stained with SiR-actin. **o,** The Pearson correlation coefficient of T7-GFP and SiR-actin on retraction fibers was calculated based on the images in **n**. N = 40 images were analyzed. Values between 0 and −0.3 indicate weak negative correlations. **p,** Snapshots from a time-lapse video of a migrating NRK cell stably expressing T7-GFP and stained with SiR-actin. The cropped area is where retraction fibers are initiated from the rear of the cell. See also **Supplementary video 3**. Scale bars, 10 μm in **b**; 5 μm in **c, h, n** and insert in **b**; 2 μm in **d** and **g**.

### TSPAN7 is immobile on tubular membranes

By serendipity, we performed fluorescence recovery after photobleaching (FRAP) assays and discovered that TSPAN7-GFP was immobile on reconstituted tubular membranes (Fig. 2d, e), an unexpected observation for such a small transmembrane protein. Accordingly, photobleaching of TSPAN7-GFP on retraction fibers in cells also resulted in no significant fluorescence recovery (Fig. 2g, i), indicating that TSPAN7-GFP remains immobile within cellular structures. Similarly, TSPAN7-GFP displayed immobility on TNTs (Fig. 2k and Extended Data Fig. 4a). Furthermore, endogenous TSPAN7 on retraction fibers in primary LECs exhibited the same immobile behavior (Fig. 2l and Extended Data Fig. 4b). Together, these results demonstrate that TSPAN7 exhibits a distinctive immobility on tubular membranes.

We next investigated whether other members of the tetraspanin family also exhibit immobility on retraction fibers. Through phylogenetic analysis, tetraspanins can be classified into multiple subgroups (Fig. 2f). We overexpressed five tetraspanins fused with GFP in NRK cells: TSPAN7, its closest homologue TSPAN6 and three representative members from other subgroups, including TSPAN33, TSPAN4 and TSPAN27. Unlike TSPAN7, TSPAN6 exhibited partial recovery at a relatively slow rate, while TSPAN33, TSPAN4 and TSPAN27 showed fast and nearly complete recovery after bleaching (Fig. 2g, i and Extended Data Fig. 3a). Interestingly, the mobility of the tested tetraspanins is negatively correlated with their enrichment on retraction fibers. TSPAN7 displayed the strongest preference for retraction fibers, followed by TSPAN6, which belongs to the same subgroup as TSPAN7. Other tetraspanins exhibited notable yet relatively weak (3–6 fold on average) preference for retraction fibers compared to flat cell membranes. (Fig. 2h, j and Extended Data Fig. 3b). The negative correlation between mobility and enrichment level of the tested tetraspanins also suggests that immobility could serve as a mechanism for sequestering TSPAN7 molecules and further enhancing their local enrichment on tubular membranes.

### The immobility of TSPAN7 on tubular membranes is independent of F-actin

It is well documented that a direct interaction with cortical actin can confine the mobility of some transmembrane proteins^34^. We therefore investigated whether the actin cytoskeleton contributes to the immobility of TSPAN7 on tubular membranes. We treated cells with Latrunculin A (LatA) to disrupt the assembly of F-actin. While inducing significant rounding of cell bodies, LatA treatment did not disrupt the integrity of established retraction fibers or the immobile state of TSPAN7-GFP on them (Fig. 2m and Extended Data Fig. 4c). This result suggested that the immobility of TSPAN7 is independent of F-actin. Indeed, in cells stained with the F-actin-specific dye SiR-actin, the fluorescent signals corresponding to TSPAN7-GFP and SiR-actin rarely colocalized on retraction fibers (Fig. 2n, o). When tracking the dynamics of these two components through time-lapse imaging (Supplementary video 3), we observed an apparent relay between SiR-actin and TSPAN7-GFP during the early stage of retraction fiber formation (Fig. 2p). These data suggest that although TSPAN7 and F-actin are spatially segregated, a temporal crosstalk may exist.

### TSPAN7 forms highly ordered spiral assemblies on retraction fibers

We next sought to explore the structural basis of TSPAN7’s immobility on tubular membranes by cryo-EM. For this purpose, we performed *in situ* cryo-EM analysis on NRK cells overexpressing either TSPAN7-GFP or TSPAN4-GFP, the latter serving as a control due to its high mobility on retraction fibers (Fig. 2g, i). Briefly, NRK cells were seeded and cultured overnight on EM grids. We captured fluorescence images of the cells to identify retraction fibers with strong green fluorescent signals. The EM grids were then rapidly frozen for cryo-EM imaging. Starting with a low-magnification cryo-EM micrograph, we correlated it with the fluorescence image to pinpoint the target retraction fibers. We then switched to high-magnification mode to collect the cryo-EM images or tilt series for cryo-electron tomography (cryo-ET) of the retraction fibers (Fig. 3a).

**Fig. 3.**
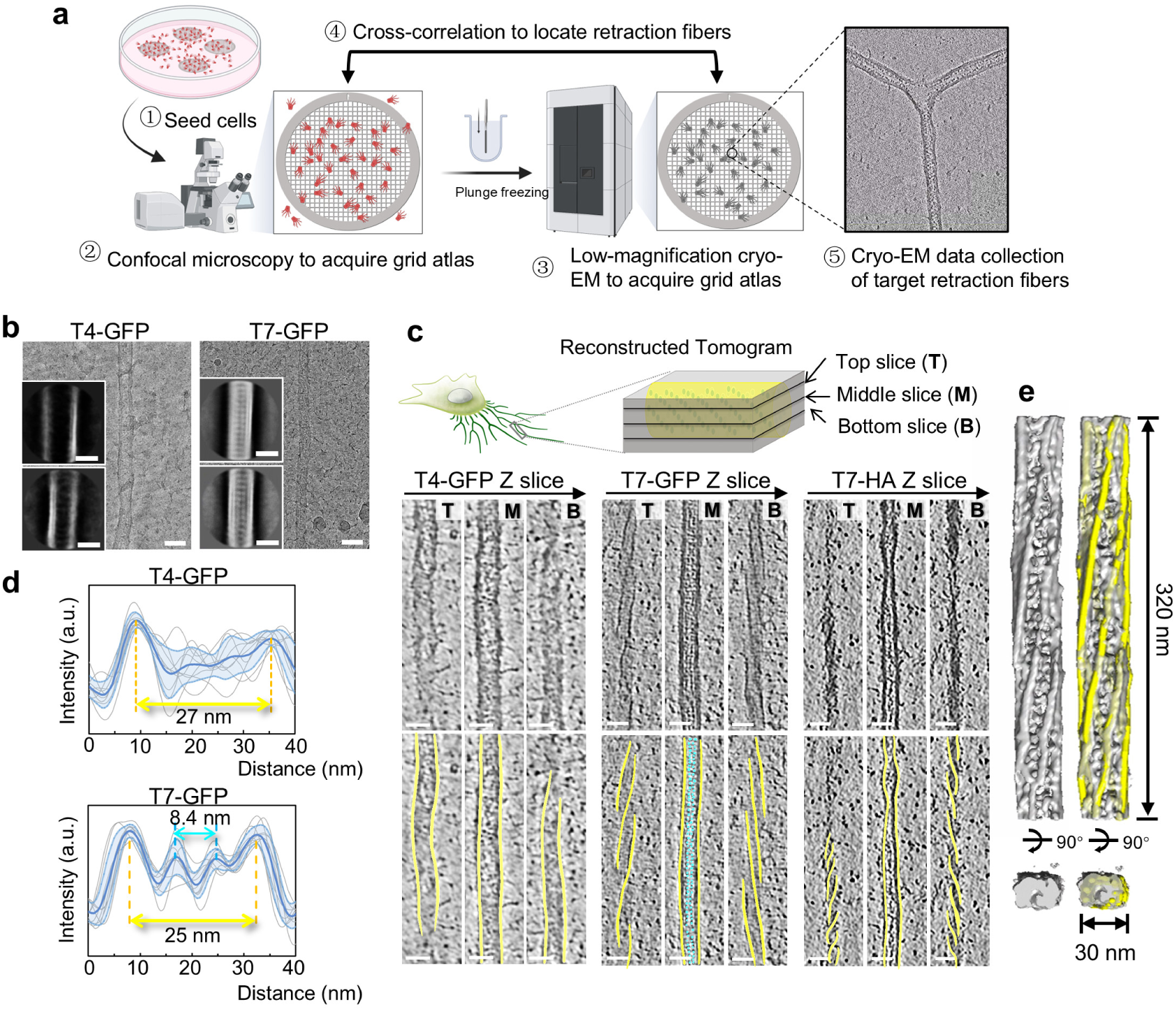
Cryo-EM reveals the highly ordered spiral assembly of TSPAN7 on retraction fibers. **a,** Diagram of cryo-EM sample preparation and data collection of *in situ* retraction fibers. First, cells were seeded on EM grids (step 1). A confocal fluorescence light microscope was used to take images of the grids (step 2). After that, the EM grids were blotted from their back side and then plunged into liquid ethane to prepare the cryo-specimens. The atlas of the EM grid was captured and cross-correlated with the fluorescence images in step 2 to locate the target retraction fibers with strong green fluorescent signals (step 4). High-magnification tilt series or single-particle datasets were then acquired at the desired regions (step 5). **b,** Two-dimensional classification results of particles picked from the single-particle cryo-EM micrographs of *in situ* retraction fibers with TSPAN4-GFP (left) or TSPAN7-GFP (right). **c,** Representative bottom (B), middle (M), and top (T) slices of the tomograms of TSPAN4-GFP (left), TSPAN7-GFP (middle) and TSPAN7-HA (right) retraction fibers. The yellow lines outline the retraction fiber boundaries in different slices, and the cyan markers indicate the density of the regular dots (GFP) on the cytoplasmic side of the cell membrane. See also **Supplementary video 4, 5, 6**. **d,** Intensity profiles of lines (N = 9) across the middle slices of tomograms of retraction fibers containing TSPAN4-GFP (upper) or TSPAN7-GFP (lower) are plotted and aligned together. The deep blue lines are the averaged traces for all plots. The light blue lines are the averaged traces with the standard deviation. Distances separating intensity peaks are indicated for retraction fiber boundary densities (yellow double-headed arrow) and luminal protein densities (blue double-headed arrow). a.u., arbitrary units. **e,** Left: Tomographic segmentation of the TSPAN7-GFP retraction fiber indicates a spiral configuration. Right: The spiral segmentation is labeled with yellow lines highlighting its six-start helical structure. Scale bars, 30 nm.

We performed both single-particle cryo-EM and cryo-ET analysis of retraction fibers with TSPAN7-GFP and found that most of them exhibited a more uniform diameter (Fig. 3b, right, and Fig. 3c, d). Notably, we observed a row of dots, each about 5 nm in size, regularly distributed along the cytoplasmic side of TSPAN7-GFP retraction fibers, a feature absent from TSPAN4-GFP retraction fibers (Fig. 3b, left, and Fig. 3c, d). We hypothesized that the observed dots corresponded to the GFP tags (27 kDa) fused to the C-terminus, i.e. the cytoplasmic side, of the TSPAN7 protein, as their size matched well with that of GFP. To test this, we replaced the GFP tag with a much smaller HA tag consisting of only nine amino acids. This substitution indeed eliminated the dot pattern on the cytoplasmic side (Fig. 3c, right). These results support the idea that the dot pattern in TSPAN7-GFP retraction fibers is associated with the GFP protein. More importantly, this pattern directly served as an indicator of the spatial organization of the fused protein, and suggested that TSPAN7 may form an ordered and packed structure in retraction fibers—a unique feature not observed with TSPAN4.

In agreement with this speculation, cryo-ET analysis revealed a spiral configuration in the TSPAN7-GFP retraction fibers (Fig. 3c, middle, Extended Data Fig. 5b and Supplementary video 4), which was absent in TSPAN4-GFP retraction fibers (Fig. 3c left, Extended Data Fig. 5a and Supplementary video 5). A similar spiral arrangement was also observed in retraction fibers containing TSPAN7-HA (Fig. 3c, right and Supplementary video 6), although these fibers were narrower than their TSPAN7-GFP counterparts. Therefore, while the diameter of TSPAN7-containing retraction fibers may be influenced by the tag size, the formation of the spiral structure is an intrinsic property of TSPAN7, independent of labeling strategies. Segmentation of the TSPAN7-GFP retraction fiber tomogram revealed a ∼30 nm-wide right-handed spiral structure with six protofilaments winding together in the tubular membrane (Fig. 3e).

We further examined the distribution and assembly of TSPAN7 spirals on retraction fibers at varying distances from the cell bodies, from proximal to distal (Extended Data Fig. 6a). In the proximal region, where membrane curvature begins to increase, no spiral architecture was observed (Extended Data Fig. 6b, upper). In contrast, a distinct spiral arrangement was clearly visible in the distal region (Extended Data Fig. 6b, middle and bottom). These results indicate that spiral assembly of TSPAN7 starts in the proximal region and becomes more ordered and stable in the distal region of the retraction fiber as it gets longer.

The spiral assembly inspired us to investigate whether TSPAN7 polymers can behave like molecular springs—capable of modulating their packing density in response to varying membrane tension levels. To investigate this, we collected cryo-EM tilt-series under three conditions with distinct membrane tension states: (1) *in situ* retraction fibers from untreated cells, (2) *in situ* retraction fibers from cells treated with LatA, which have significantly reduced membrane tension^36^, and (3) retraction fibers isolated from cells, to eliminate the lateral transfer of membrane tension from the cell body (Extended Data Fig. 7).

We first quantified the percentage of retraction fibers exhibiting clear spiral structures for each condition. Fibers from LatA-treated cells showed significantly higher spiral prevalence (88%) compared to fibers from untreated control cells (74%). Strikingly, nearly all isolated retraction fibers (98%) displayed a distinct spiral structure (Extended Data Fig. 8). We also assessed the tightness of the spiral structure using an angular velocity metric, defined as the ratio of the twist angle to the rise of the spiral (Extended Data Fig. 9). The angular velocity of TSPAN7 spirals showed a moderate yet noticeable increase in the LatA-treated sample and a more pronounced increase in the isolated sample, compared to the untreated sample (Extended Data Fig. 9). These findings show that TSPAN7 spirals are capable of maintaining their structural integrity after membrane tension release and adjusting their packing density in response to tension fluctuations.

### Molecular basis of the TSPAN7 spiral

We next sought to elucidate the molecular basis of the TSPAN7 spiral on retraction fibers. Single-particle cryo-EM analysis revealed ordered TSPAN7-GFP arrays along the retraction fibers; however, the resolution was insufficient for detailed structural insights (Fig. 4a). We performed subtomogram averaging analysis on the cryo-ET reconstructions of the ordered TSPAN7-GFP spirals on retraction fibers and obtained a rather low-resolution 3D map of a six-start spiral architecture, with GFP density localized at the center of the spiral (Fig. 4b, Extended Data Fig. 10, and Supplementary Table 1). The limited resolution in these reconstructions is likely due to: (1) heterogeneity in spiral assembly, (2) flexibility introduced by the GFP tag, and (3) the small size of the TSPAN7 protein itself, which impedes precise particle alignment. Despite the limited resolution, the above reconstructions suggested that a dimer of TSPAN7 likely serves as the building unit of the spiral assembly (Fig. 4a-b).

**Fig. 4.**
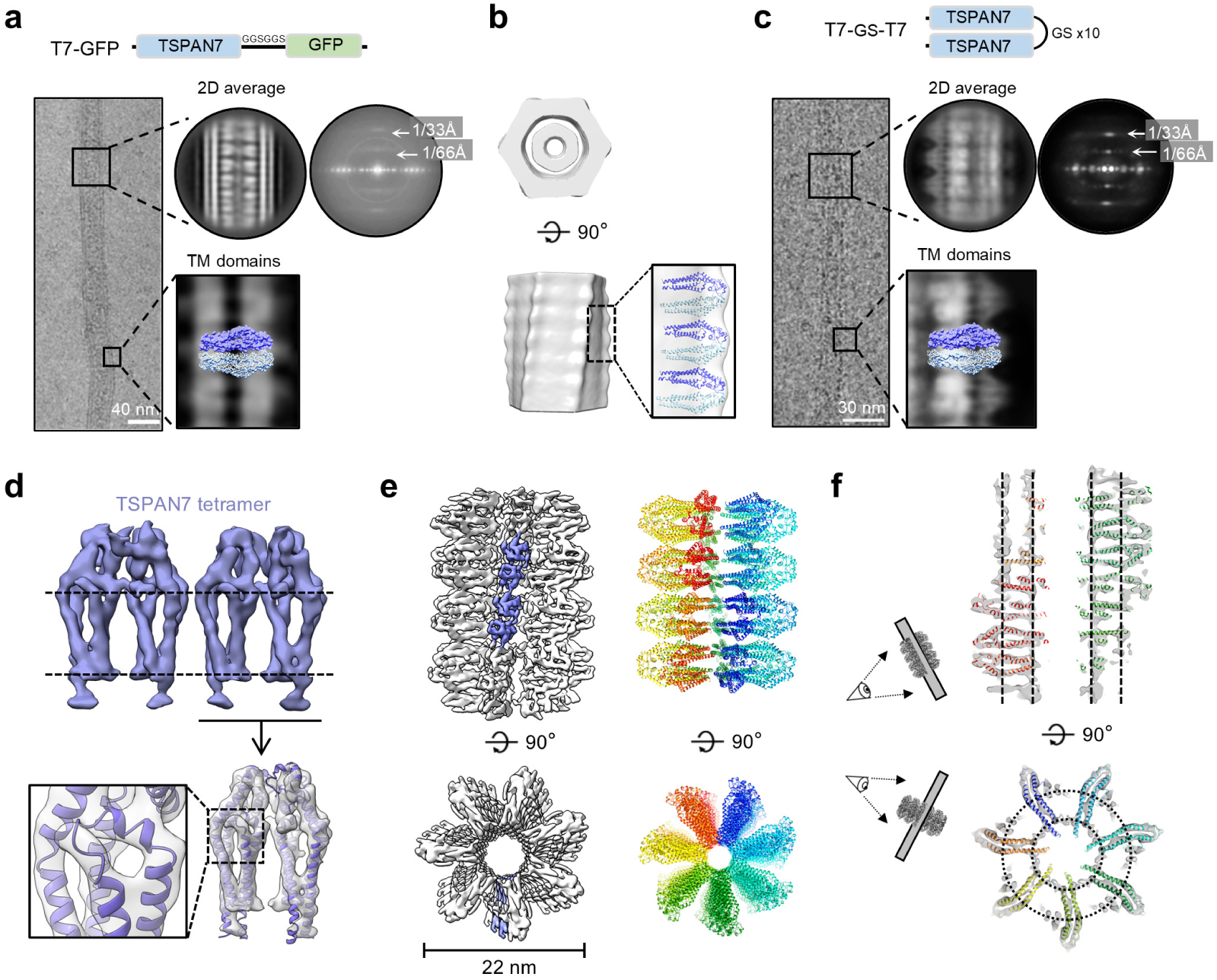
Cryo-EM reconstruction of the TSPAN7 spiral. **a,** Cryo-EM analysis of retraction fibers containing TSPAN7-GFP (T7-GFP). Left: Cryo-EM micrograph of a retraction fiber. Right: Representative 2D class average of retraction fiber particles with its Fourier transform image showing layer line indexing (top) and transmembrane (TM) domains with a superimposed TSPAN7 dimer (bottom). **b,** Subtomogram averaging result of retraction fibers containing TSPAN7-GFP. The enlargement of the boxed region shows that a single TSPAN7 dimer nicely fits each repeating unit of the spiral structure. **c,** Cryo-EM analysis of retraction fibers containing engineered TSPAN7 dimers (T7-10×GS-T7). **d-e,** Cryo-EM reconstruction of the locally refined TSPAN7 tetramer (**d**) and the TSPAN7 spiral (**e**). Dashed horizontal lines in **d** (top) indicate the plasma membrane. **f,** Cross-sections of the TSPAN7 spiral structure with the docked model, showing nice density fitting.

Based on these observations, we engineered a TSPAN7 dimer construct by covalently linking two monomers and removing the flexible GFP in order to improve sample homogeneity for high-resolution reconstruction. NRK cells overexpressing this TSPAN7 dimer construct produced retraction fibers with similar morphology and yield to those with TSPAN7-GFP (Extended Data Fig. 11a-b). Cryo-ET analysis revealed more uniform spiral architectures in the TSPAN7-dimer-expressing retraction fibers (Extended Data Fig. 11c). Notably, single-particle cryo-EM analysis demonstrated significantly enhanced structural details in the dimer-containing fibers compared to TSPAN7-GFP fibers, indicating a high homogeneity of the helical assembly. Layer-line indexing of the Fourier transforms of TSPAN7-GFP retraction fiber class averages and TSPAN7-dimer retraction fiber class averages confirmed that both constructs maintain the same helical rise parameter (Fig. 4a, c).

Using cryo-EM datasets of retraction fibers containing the engineered TSPAN7 dimers, we performed single-particle analysis and obtained a 7.8 Å resolution reconstruction of the entire spiral architecture, with local refinement of the TSPAN7 density reaching 5.9 Å resolution (Fig. 4d, e, Extended Data Fig. 12, and Supplementary Table 2). The cryo-EM maps revealed discernible secondary structure features of TSPAN7. By integrating these densities with AlphaFold3 predictions^37^, we built models of both the TSPAN7 tetramer in a protofilament and the complete spiral assembly in retraction fibers (Fig. 4d-f, Extended Data Fig. 12). The reconstruction and model of retraction fibers containing the engineered TSPAN7-dimer spiral revealed that seven TSPAN7 protofilaments assemble into a right-handed spiral with a diameter of 22 nm that spans the membrane tubule. Within each protofilament, TSPAN7 protomers organize in head-to-head and back-to-back configurations via two distinct interaction interfaces. We also observed that a small percentage of spirals were composed of six or eight protofilaments (Extended Data Fig. 13), indicating a certain degree of flexibility in the assembly process.

We examined the TSPAN7 spiral model built based on the 3D cryo-EM map and revealed two critical interaction interfaces (IFs) involved in the formation of spiral assemblies (Fig. 5a): (1) one centered on glutamate 115 (E115; Fig. 5b) and (2) the other consisting of a highly charged short α-helix (Helix 2) in the large extracellular loop (aa 135-147; Fig. 5c). To investigate the functional role of these sites, we generated NRK cells expressing TSPAN7-GFP with a double mutation (E115A/ΔHelix2) and then collected cryo-EM tilt series of the retraction fibers with strong green fluorescent signals. Cryo-ET analysis revealed abundant GFP density within the retraction fibers but a complete absence of TSPAN7 spiral organization (Fig. 5d, Supplementary video 7). These results demonstrated that while mutations at these two interfaces preserved TSPAN7’s ability to localize in retraction fibers, they abolished its capacity to form spiral assemblies.

**Fig. 5.**
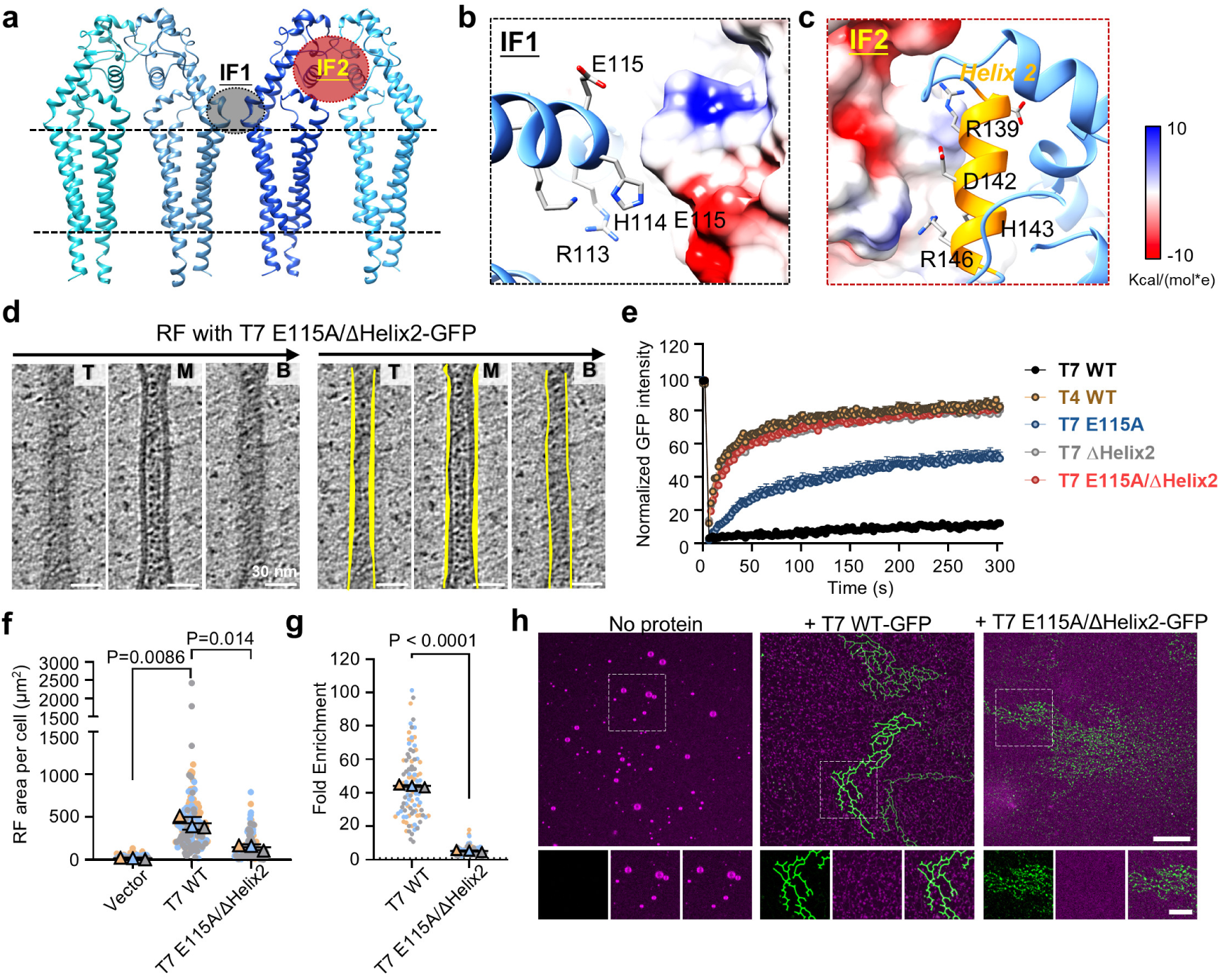
Molecular mechanism of assembly of the TSPAN7 spiral. **a,** TSPAN7 tetramer model, highlighting two interaction interfaces between protomers (IF1 and IF2). **b-c,** Close-up views of IF1 (**b**) and IF2 (**c**), with one protomer shown as a ribbon and the other shown as an electrostatic potential surface. Key residues mediating the interactions are labeled. **d,** Left: Representative tomogram slices (top, T; middle, M; bottom, B) of a retraction fiber (RF) containing the TSPAN7 double mutant (E115A/ΔHelix2, see Supplementary Video 7). Right: Same slices with boundaries outlined in yellow. **e,** FRAP analysis of retraction fibers from NRK cells expressing wild-type TSPAN7-GFP (T7 WT), wild-type TSPAN4-GFP (T4 WT), TSPAN7 single mutant (T7 E115A or T7 ΔHelix2), or the TSPAN7 double mutant (T7 E115A/ΔHelix2). Curves represent normalized mean + SEM from 15 bleached regions. See also **Extended Data Fig. 15a**. **f,** Quantification of RF area in NRK cells expressing GFP (vector), T7 WT, or T7 E115A/ΔHelix2. Cell numbers analyzed: GFP (N = 154), T7 WT (N = 156), T7 E115A/ΔHelix2 (N = 155). See also **Extended Data Fig. 15b**. **g,** The fold enrichment 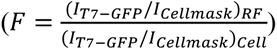 of GFP intensity on RFs versus cell bodies in NRK cells expressing T7 WT (N = 96) or T7 E115A/ΔHelix2 (N = 98). See also **Extended Data Fig. 15c**. **h**, Representative confocal images showing the morphology of reconstitution systems without protein (left), with T7 WT-GFP (middle panel) or with T7 E115A/ΔHelix2-GFP (right panel) after leaving at room temperature overnight. Scale bars, 30 nm in **d**; 5 μm in **h**.

FRAP analysis of TSPAN7 on retraction fibers revealed that TSPAN7-GFP with either E115A or ΔHelix2 mutations exhibited partial loss of immobility, with the ΔHelix2 mutant showing greater mobility. The double mutant (E115A/ΔHelix2) displayed nearly complete loss of immobility, approaching TSPAN4-like mobility levels (Fig. 5e, Extended Data Fig. 15a). The double mutant also exhibited a diminished capacity to promote retraction fiber formation (Fig. 5f, Extended Data Fig. 15b), as well as reduced enrichment on retraction fibers (Fig. 5g, Extended Data Fig. 15c). Finally, we performed the *in vitro* reconstitution assay (described in Fig. 2a) using recombinant wild-type or double mutant TSPAN7. While GUVs embedded with wild-type TSPAN7 maintained tubular networks after overnight incubation (Fig. 5h, middle), those embedded with the double mutant TSPAN7 failed to produce tubular structures with prolonged stability (Fig. 5h, right). These observations indicate that TSPAN7 spiral assembly is critical for sustaining the structural integrity of reconstituted membrane tubules.

Moreover, sequence alignment revealed that the identified interaction sites are fully conserved in mouse, rat and human TSPAN7 (Extended Data Fig. 14a). Alignment of human tetraspanins revealed that E115 is conserved in TSPAN6 but not in TSPAN4, while the sequence of Helix 2 is not conserved in TSPAN6 or TSPAN4 (Extended Data Fig. 14b).

Collectively, these data indicate that the identified interaction interfaces are crucial for TSPAN7’s oligomerization and the TSPAN7 spiral model presented above faithfully represents how TSPAN7 protomers organize into their spiral configuration on tubular membranes. Our results also highlight the importance of spiral assembly in shaping and stabilizing tubular membranes.

### TSPAN7 spirals stabilize tubular membranes under shear stress

Shear stress, which arises from fluid flow or mechanical forces, can induce membrane deformation by exerting tangential forces on the lipid bilayer. In the case of tubular protrusions, these forces can lead to their elongation and narrowing. To further investigate the mechanism by which TSPAN7 spirals stabilize tubular membranes, we established a flow assay where cells were cultured in a microscopy-compatible channel device and a controlled, unidirectional shear flow was applied. The deformation of tubular protrusions on the cell surface in response to shear stress was captured and analyzed quantitatively (Fig. 6a). It should be noted that, unlike the biogenesis of retraction fibers—which involves the incorporation of newly synthesized proteins and typically takes hours for a significant extension—the elongation of tubular protrusions under shear flow occurs in minutes. This rapid response leaves little room for the involvement of newly synthesized proteins. Therefore, the assay is particularly suited for evaluating the deformation of established tubular protrusions, rather than the *de novo* formation of such structures in cells. This approach allows us to focus on the mechanical and structural aspects of tubular membrane stability under dynamic conditions.

**Fig. 6.**
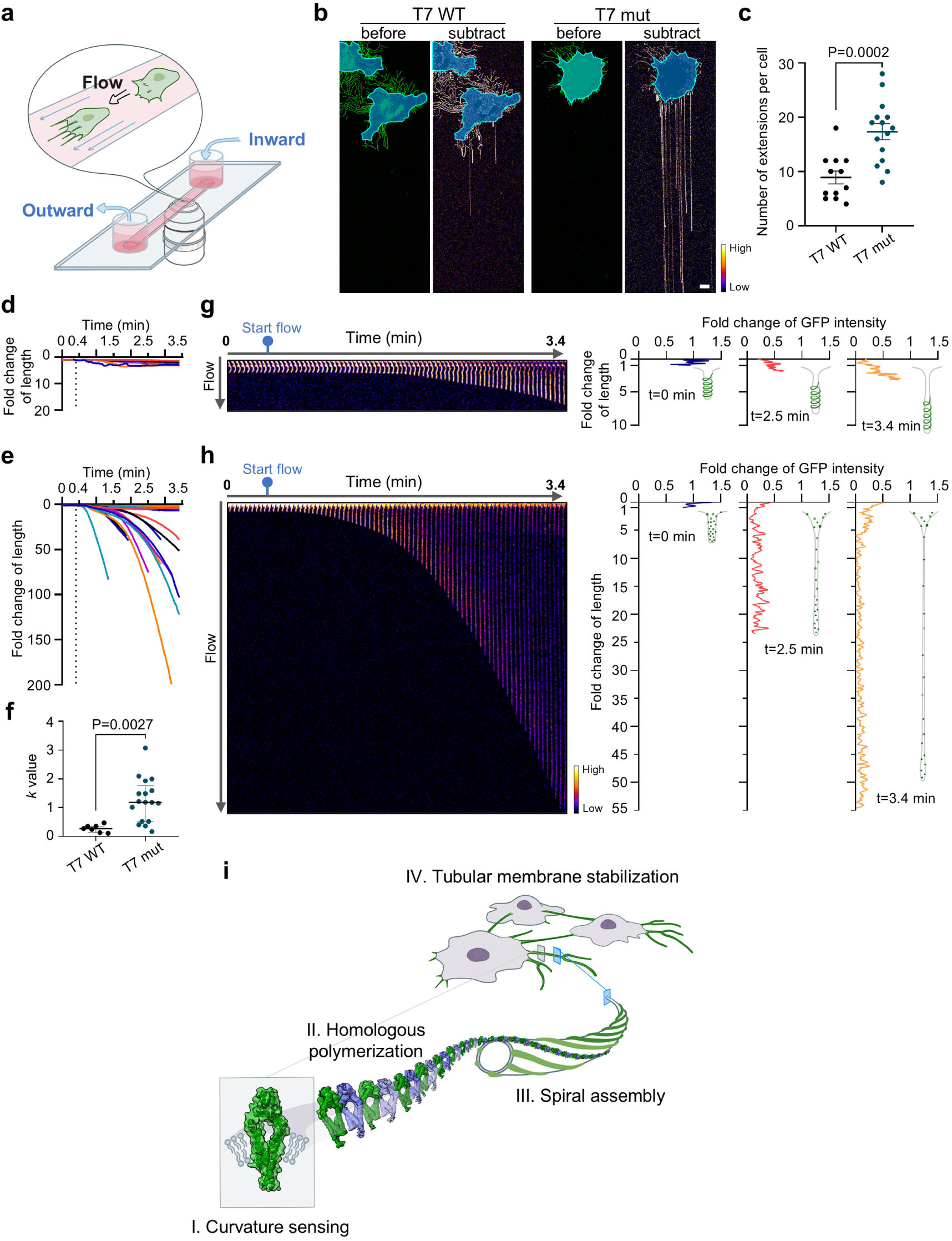
TSPAN7 spirals stabilize tubular membranes under shear stress. **a,** Schematic illustration of the flow assay. Cells cultured in a narrow channel were exposed to a controlled, unidirectional flow of PBS and the progress of membrane deformation was captured by time-lapse imaging. **b-h,** NRK cells stably expressing wild-type TSPAN7-GFP (T7 WT) or TSPAN7 E115A/ΔHelix2-GFP (T7 mut) were exposed to a 3 min course of PBS flow. The shear stress was 21.83 dyn/cm^2^. Subtracted images created by *I_substract_* = *I_after flow_* - *I_Before flow_* are presented as heatmaps in b. The areas corresponding to cell bodies are colored blue (See also **Supplementary video 8**). The number of tubular extensions per cell was analyzed and plotted in **c**. N = 12 and 15 cells were analyzed for T7 WT and T7 mut, respectively. A representative cell from each group was selected for further analysis (See also **Supplementary video 8)**. The fold change of the length of tubular protrusions over time is plotted in **d** (T7 WT) and **e** (T7 mut); each curve represents a tubular protrusion. N = 10 and 23 tubular protrusions were analyzed for T7 WT and T7 mut, respectively. Non-linear regression was performed on these curves and the *k* values with R square values above 0.75 are shown in **f**. A representative tubular protrusion from each group was selected for further analysis in **g** (T7 WT) and **h** (T7 mut). Kymographs of tubular protrusion extension under flow are shown in the left panels, and the distributions of GFP intensity along the tubular protrusions at the indicated time points (t=0, 2.5, 3.4 min) are shown in the right panels. The inset illustrations are models of protein assembly on tubular protrusions. **i,** Working model of the transmembrane skeleton formed by TSPAN7. TSPAN7 monomers sense high curvature, forming polymers via homologous interaction on tubular membranes. The polymer chains shape and stabilize the tubular membranes through spiral assembly. Scale bars, 10 μm in **b**.

We applied a 3-minute course of shear flow, with a strength of 21.83 dyn/cm^2^, to cells expressing either the spiral-forming wild-type TSPAN7-GFP (wild-type TSPAN7) or the spiral-deficient TSPAN7-GFP double mutant (mutated TSPAN7) (Fig. 5d). The elongation of membrane protrusions was abundant and significant in cells expressing mutated TSPAN7 (Fig. 6b, c and Supplementary video 8). In sharp contrast, only a small population of membrane protrusions showed detectable elongation in cells expressing wild-type TSPAN7 (Fig. 6b, c and Supplementary video 8).

To analyze the kinetics of membrane deformation in detail, we tracked individual tubular protrusions in representative cells and plotted the fold change in length as a function of time (Fig. 6d, e and Supplementary video 8). While the majority of tubular protrusions elongated extensively (∼40 fold) in cells expressing mutated TSPAN7 (Fig. 6e), they showed a limited degree (less than four fold) of elongation in cells expressing wild-type TSPAN7 (Fig. 6d). We then performed non-linear regression on these curves using the exponential growth equation (*Y* = *Y*_0_*e^kx^*) and the best-fit *k* values were calculated (Fig. 6f). The results showed that the deformation of tubular protrusions in response to a given force progressed much faster in cells expressing mutated TSPAN7 than in cells expressing wild-type TSPAN7.

Furthermore, we monitored the distribution of the fluorescence intensity of wild-type TSPAN7 and mutated TSPAN7 on these tubular protrusions (Fig. 6g, h). We found that the fluorescence intensity of the mutated TSPAN7 decayed significantly and successively along the elongated tubular protrusion, indicating the extreme narrowing of the tube (Fig. 6h). In contrast, the fluorescence intensity of wild-type TSPAN7 showed a smaller reduction, and a strong continuous TSPAN7 signal was detected in the protrusion. As the protrusion elongated, the strong continuous TSPAN7 signal was confined to the distal part, with a much weaker signal in the more proximal part of the tube (Fig. 6g). This result indicates that extreme tube narrowing is effectively prevented in regions where TSPAN7 spirals exist. Collectively, these results highlight the contribution of TSPAN7 spiral assembly in resisting membrane deformation in response to mechanical stress, through the effective restriction of tube narrowing.

In summary, we have identified TSPAN7 as a curvature-sensitive protein that spontaneously concentrates and polymerizes into a spiral configuration on highly curved tubular membranes. This spiral assembly markedly enhances the formation of membrane tubules and substantially improves their stability (Fig. 6i). Given the structural parallels between TSPAN7 tubules and microtubules—both composed of polymerized basic protein units forming linear protofilaments and both providing structural support—we suggest that TSPAN7 tubules act as a form of “transmembrane skeleton”. This transmembrane skeleton, akin to the cytoskeleton, not only offers structural support but may also fulfill various other roles.

## Methods

### Reagents and antibodies

Fibronectin was purchased from Invitrogen (PHE0023) or Sigma (F0895). WGA-TMR (Wheat Germ Agglutinin tetramethylrhodamine conjugate, W849) and CellMask Deep Red (C10046) were purchased from Invitrogen. SiR-Actin Kit (CY-SC001) was purchased from Cytoskeleton. Latrunculin A (10010630) was purchased from Cayman. PEI MW40000 was purchased from Yeasen. Lipids including DOPC, DOPE, POPS, cholesterol, 16:0 Biotinyl Cap PE and 18:1 Liss Rhod PE were purchased from Avanti. Streptavidin (85878) was purchased from Sigma. Polybead® Polystyrene Sampler Kit (19822-1) was purchased from Polysciences.

Antibodies used in western-blot analysis include anti-GFP (Roche, 11814460001), anti-GAPDH (Proteintech, 60004-1-Ig), anti-integrin α5 (CST, 4705T), anti-calnexin (Abcam, ab22595), anti-Lamp2 (Sigma), anti-Tim23 (BD, 611222). Antibodies used in flow cytometry analysis include Alexa Fluor 594 anti-gp38 (BioLegend, 156205), Alexa Fluor 647 anti-CD31 (BioLegend, 102516), Alexa Fluor 700 anti-CD45 (eBioscience, 56-0451-82), PE anti-TER-119 (BioLegend, 116208).

### Plasmid construction

For transient transfection and imaging, the coding sequences of TSPAN7 and other tetraspanins were amplified from rat cDNA through polymerase chain reaction (PCR) and subsequently cloned into a pEGFP-N1 vector (linearized with XhoI and BamHI) through homologous recombination. TSPAN7 constructs with point mutations were generated using a standard site-directed mutagenesis protocol^1^. TSPAN7 constructs with deletions were generated using a standard overlap extension PCR protocol^2^.

For protein purification, the coding sequence of TSPAN7 was amplified from human cDNA and cloned into a modified pCAG vector. Briefly, an affinity cassette containing two repeats of Strep-Tag II followed by one FLAG tag was fused to the N-terminus of TSPAN7, and an EGFP tag was fused to the C-terminus of TSPAN7 using the linker GGSGGS.

For the construction of TSPAN7 dimer plasmids, two full-length TSPAN7 proteins were linked through ten repetitive GS sequences and inserted into a pCAG vector containing two repeats of Strep-Tag II.

### Cell culture and transfection

NRK (ATCC CRL-6509) cells were cultured at 37 °C and 5% CO_2_ in DMEM high glucose medium supplemented with 10% fetal bovine serum, 1% penicillin-streptomycin and 1% Glutamax. HEK293T (ATCC CRL-3216^TM^) cells were cultured at 37 ° C and 5% CO_2_ in DMEM (Gibco-Thermo Fisher scientific 11995065) supplemented with 10% fetal bovine serum (Biological Industries, 04-001-1A) and 1% penicillin-streptomycin. HEK293F cells were cultured in suspension at 37 °C, 5% CO_2_ and 120 rpm in SMM293-TII supplemented with 1% penicillin–streptomycin. Primary cells isolated from murine lymph nodes were cultured at 37 °C and 5% CO_2_ in complete Endothelial Cell Medium (SC1001). Electroporation (Amaxa nucleofector, program X-001) was used for the transfection of NRK cells. PEI was used for the transfection of HEK293F cells.

### Mice

WT C57BL/6 (Jax 000664)-specific pathogen-free (SPF) mice were purchased from the Laboratory Animal Center of Tsinghua University, China. The TSPAN7-GFP knock-in mouse strain was constructed by Cyagen Biosciences. Genotyping of the knock-in mice was performed using the following primer pairs:

Null-F 5’-GCAGGGGATAGCTTTGTGACG-3’

Null-R 5’-ACCTCTTCCCCAACATACGG-3’

KI-F 5’-GCAGGGGATAGCTTTGTGACG-3’

KI-R 5’-CGTAGGTGGCATCGCCCTC-3’

All mice were bred and kept under SPF conditions in the Laboratory Animal Center of Tsinghua University in accordance with the National Institutes of Health Guide for Care and Use of Laboratory Animals.

### Isolation of GFP^+^ cells from mouse lymph nodes

6 to 8-week-old male TSPAN7-GFP knock-in mice were used for the isolation. For microscopic experiments, the isolation procedure was modified from published protocols^3^. Briefly, mice were anesthetized by intraperitoneal injection of Avertin. Inguinal, mesenteric and axillary lymph nodes were carefully dissected, minced with fine scissors and digested with an enzyme cocktail (0.1% (w/v) collagenase II, 0.25% (w/v) collagenase IV, 15 mg/mL DNAse I, 4 mM CaCl_2_ and 4% (v/v) heat-inactivated FBS in DMEM) at 37°C for 30 min. The resulting cell suspension was filtered by passing through a 70-μm cell strainer, centrifuged at 300 x g for 5 min and then resuspended with ECM. GFP^+^ cells were then sorted by flow cytometry (Beckman Coulter, MoFlo Astrios EQ).

### Characterization of GFP^+^ cells from murine lymph node

6 to 8-week-old male wildtype C57BL/6 mice and TSPAN7-GFP knock-in mice were used for the assay. Single-cell suspensions of lymph nodes were prepared and stained following a published protocol^4^. The staining antibody cocktail was slightly modified (gp38-AF594, CD31-AF647, CD45-AF700 and Ter119-PE). The sample was then washed, resuspended with FACS buffer containing DAPI and analyzed by flow cytometry (Beckman Coulter, CytoFLEX LX).

### Tether pulling from immobilized GPMVs

NRK cells at 90% confluence were washed with GPMV buffer (10 mM HEPES, 2 mM CaCl_2_, 150 mM NaCl, pH 7.4) and incubated with vesiculation buffer (GPMV buffer supplemented with 2 mM DTT and 25 mM formaldehyde before use) for 1 h at 37 °C. The supernatant containing GPMVs was collected and incubated with 10 μg/ml biotin-PE (and 0.25 μg/ml Rhodamine-PE in some cases) for 10 min at room temperature. Polystyrene beads (2 μm diameter) were incubated with 100 μg/ml streptavidin for 1 h. GPMVs were added to a plasma-treated glass surface and settled for 1 h before the addition of polystyrene beads. Optical tweezer experiments were performed on a Nikon HD25 confocal microscope equipped with an MMI CellManipulator. A polystyrene bead was captured by a laser trap and was allowed to contact with a GPMV. By pulling the bead in the opposite direction, a membrane tether was generated. Z-stack images covering the GPMV and the membrane tether were collected using an A1 confocal microscope.

### Tether pulling from aspirated GPMVs

HEK293T cells were used to generate GPMVs. The experiments were performed using a C-trap® confocal fluorescence optical tweezers setup (LUMICKS, Amsterdam, the Netherlands) made of an inverted microscope based on a water-immersion objective (NA 1.2) together with a condenser top lens placed above the sample. The methods regarding GPMV preparation, experimental setup and data analysis were described in detail before^5^. Briefly, a membrane tube was pulled from aspirated GPMVs using beads trapped by the optical tweezers. First, a membrane tube was pulled at relatively low suction pressure (0.05-0.1 mbar). Then the suction pressure was gradually increased (usually by 0.05-0.1 mbar steps) until values in the range of 0.4-2.5 mbar were reached.

### Protein expression and purification

HEK293F cells were grown to a density of 0.8-1.2 × 10^6^ cells/mL and then transfected with 2 mg plasmid per liter of cell culture using PEI (DNA: PEI = 1: 2.5). 48-72 hours after transfection, cells were harvested by centrifugation at 550 × g. The pellet was washed once with PBS and resuspended in Tris buffer (100 mM Tris, 150 mM NaCl, pH 8.0) supplemented with protease inhibitor. Cells were lysed by grinding 100 times using a glass Dounce homogenizer on ice. The lysate was centrifuged at 30,000 × g for 1 h and the pellet was resuspended with 10 mL Tris buffer supplemented with protease inhibitor. To extract TSPAN7-GFP protein from membranes, 2.5 mL of 10 % (w/v) Fos-Choline-12 was added. The mixture was rotated at 4 °C for 2 h and then centrifuged at 12,000 × g for 20 min. The supernatant containing extracted protein was loaded onto a column containing 1 mL pre-balanced strep-tactin beads and incubated for 1 h at 4 °C on a rotator. The beads were washed with 10 mL wash buffer (100 mM Tris, 150 mM NaCl, pH 8.0, containing 0.1 % (w/v) Fos-Choline-12). Protein was eluted with 8 mL elution buffer (100 mM Tris, 150 mM NaCl, 10 mM desthiobiotin, pH 8.0, containing 0.1 % (w/v) Fos-Choline-12) and concentrated using a 30 kDa MWCO centrifugal filter.

### Liposome preparation

DOPC, DOPE, POPS and cholesterol were mixed at a ratio of 4:2.5:2.5:1, making up 3 mg lipids in total. 0.025 % rhodamine-PE was added as a fluorescent indicator. The lipid mixture (prepared in a capped glass vial) was gently dried under a stream of nitrogen and left for 1 h at 37 °C to ensure complete solvent evaporation. The resulting lipid film was hydrated by adding 300 μL HEPES buffer (20 mM HEPES, 150 mM NaCl, pH 7.4) and shaking vigorously at 37 °C for 1 h. The mixture containing liposomes was transferred into a centrifuge tube and freeze-thawed for at least 10 times by transferring between liquid nitrogen and a 42°C water bath. To homogenize the liposomes, a polycarbonate membrane with 1 μm pore size was inserted into an extruder and liposomes were passed through for a total of 21 times.

### Proteoliposome preparation

Liposomes were kept at 4 °C on a rotator and protected from light during the whole process. First, liposomes (prepared as described in the previous section) were incubated with 0.1 % Fos-Choline-12 for 30 min. Then, purified TSPAN7-GFP protein was added at a protein/lipid ratio of 0.01 (for protein-free liposomes, an equal amount of wash buffer (100 mM Tris, 150 mM NaCl, 0.1 % (w/v) Fos-Choline-12, pH 8.0) was added). The mixture was incubated for 1 h. To remove the detergent, 10–12 mg of bio-beads / 100 μL liposomes were added every hour for a total of 3 times. After the third addition of bio-beads, the mixture was stirred overnight. A fourth portion of bio-beads was added on the next day, followed by 1 h incubation. The supernatant was transferred into a clean centrifuge tube and stored in aliquots at −80°C.

### GUV preparation and reconstitution

GUVs were prepared via the electroformation method using a Vesicle Prep-Pro system (Nanion). Before use, ITO (indium tin oxide)-covered glass slides were gently cleaned by liquid soap, triple-washed using distilled water/ethanol, and dried under a stream of nitrogen. A rubber ring was stuck onto the ITO-covered side of the slide, creating a confined chamber. 20 μL of proteoliposomes were deposited onto the confined glass surface and air-dried. The chamber was gently filled with 280 µL of 300 mM sorbitol and electroformation was then carried out at 0.24 V, 9.9 Hz for 90 min at 37°C. The solution containing GUVs was collected and stored at 4 °C before use.

For *in vitro* reconstitution, GUVs were added to a glass-bottom confocal chamber treated with plasma cleanser. The chamber was sealed by parafilm and incubated for at least 1.5 h before imaging by a confocal microscope.

### Imaging and staining

Cells were seeded on a 10 μg/mL fibronectin-coated confocal chamber (Cellvis) and transferred to a Nikon A1 HD25 confocal microscope equipped with a live-cell imaging system. Images were acquired using a 100× objective at 1024 × 1024 pixels. For long-term time-lapse experiments, cells were allowed to migrate for about 6 h before imaging. Images were acquired at 2 min intervals for the indicated time periods. For confocal snapshot images and FRAP assays, cells were allowed to migrate for about 16 h before imaging. For the observation of TNTs, Z-stack images were acquired. An additional staining step was performed before imaging in some experiments. Specifically, cells were incubated with 1 μg/mL WGA-TMR for 10 min, 1.25 μg/mL CellMask Deep Red for 10 min, or 1 μM SiR-actin for 1 h before imaging.

### FRAP assay and data processing

For each cell analyzed, 7 ROIs (Regions Of Interest) were defined: 3 stimulation ROIs (1.6 μm diameter), 3 reference ROIs (1.6 μm diameter) and 1 background ROI. The experiment contains three phases: first, two frames were captured before stimulation; next, stimulation ROIs were bleached with 100% 488 nm laser for 1.5 s; finally, a 5 min time-lapse movie was recorded after bleaching. Data processing was performed using the NIS-Elements software. First, alignment was performed to correct XY motion. Then, the mean fluorescent signals of 7 ROIs over time were measured. Next, the fluorescent signals of stimulation ROIs were subjected to background correction and reference correction. Finally, all datasets were linearly normalized to the [0, 100] range. The largest value of each dataset was defined as 100% and 0 was defined as 0%.

### Isolation of retraction fibers from cell culture for cryo-EM

NRK cells stably expressing TSPAN7-GFP or the engineered TSPAN7 dimer were seeded into 15-cm culture dishes that were pre-coated with 4 μg/mL fibronectin and cultured overnight. Typically, 30 dishes were used for each preparation. In order to remove the majority of cell bodies while keeping retraction fibers attached to the dish, cells were washed twice with PBS, treated with 5 mM EGTA at 37 °C for 30 min, and then gently detached with a 10 mL pipette. The supernatant containing cell bodies was discarded and the culture dishes were washed twice with PBS. The remaining retraction fibers in the dish were subjected to mild trypsin treatment (0.025% trypsin-EDTA) for 3 min at room temperature and then collected into a 50 mL conical centrifuge tube. In order to remove cell debris, the sample was sequentially centrifuged at 300 × g, 500 × g, and then passed through a gravity-driven parylene filter with 6-μm pore size. Retraction fibers were then pelleted by 20,000 × g centrifugation.

### Flow assay and data processing

Channel slides (Ibidi µ-Slide VI ^0.1^ (Ibidi, 80666)) and culture medium were placed into the incubator for equilibration the day before seeding the cells. A 1 mL sterile syringe with a Luer tip was used to fill the channels with coating solution or cell suspension. Before seeding cells, channels were coated with 0.3 mg/mL fibronectin for 1 h. Remaining fibronectin was removed by perfusing the channels with culture medium. The trypsinized cell suspension was diluted to 2 × 10^5^ cells/mL and added into the channels; any extra volume of cell suspension in the reservoirs was aspirated. The µ-Slide was settled for 10 min in the incubator, allowing the attachment of cells. Afterwards, each reservoir was filled with 60 μL culture medium, and the cells were cultured for 16 h.

The flow assays were performed on a Nikon A1 Rsi HD25 confocal microscope equipped with a live-cell imaging system. The µ-Slide was connected to a microinjection pump (Longer Pump, model LSP01-2A) for perfusion. The flow rate was set to 300 μL/min, corresponding to 21.83 dyn/cm^2^ in shear stress. The calculation was performed based on the following equation in the manufacturer’s instruction:

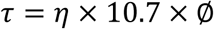

where *τ*(*dyn*/*cm*^2^) is shear stress, *η*(*dyn* · s/*cm*^2^) is dynamic viscosity, Ø(*μl*/*min*) is flow rate, and 10.7 is a slide-dependent factor. *η* = (*dyn* · s/*cm*^2^) which is the dynamic viscosity of water at 37°C, was used for calculation.

Time-lapse images (1024 × 1024 pixels) were acquired with no delay using a 60× objective. First, ten frames were captured before perfusion; next, a 3 min PBS flow was applied during imaging.

The tracking of tubular protrusion extension over time was performed manually in ImageJ. To calculate the fold change, the length of tubular protrusion in each frame was normalized to the average length of tubular protrusions in the first ten frames. Plotting and non-linear regression of the curves were performed in GraphPad.

### Scanning electron microscopy

Silicon wafers were coated with 10 µg/mL fibronectin for 1 h. Cells were then seeded on silicon wafers and cultured for 16 h before fixation with 2.5% glutaraldehyde. After washing with 0.1 M PB buffer (2.579g Na_2_HPO_4_•12H_2_O, 0.437g NaH_2_PO_4_•2H_2_O in 100 ml distilled water (pH 7.2)), samples were washed in PB buffer for a total of 4 times (15 min each), then incubated with 1% osmium tetroxide/1.5% potassium ferricyanide for 60 min at room temperature. After washing in distilled water, samples were dehydrated in an ethanol series (50, 70, 80, 90, 100, 100 and 100%; 2 min each). The samples were dried in a critical point dryer (Leica EM CPD300; Leica Microsystems). A 10-nm gold layer was sputtered onto the surface of the samples. Images were then acquired using an FEI Helios NanoLab G3 UC scanning electron microscope.

### Cryo-EM specimen preparation

For the preparation of *in situ* cryo-EM specimens, Quantifoil R1.2/1.3 Au-grids (200 mesh) were coated with homemade graphene film^6^ and glow-discharged for 15 seconds (Harrick Plasma, low-power mode). Afterwards, the EM grids were placed in a 35-mm Ibidi μ-Dish under ultraviolet illumination for 10 minutes. Next, human plasma fibronectin (FC010, Millipore, USA) solution (10 μg/mL in PBS buffer) was added onto the EM grids and incubated for 40 minutes at 37 °C. The fibronectin solution was then removed, and 1.2 × 10^5^ NRK cells were seeded onto the EM grids and cultured in an incubator overnight at 37°C and 5% CO_2_ atmosphere. After a 14-16 h culture, the EM grids were imaged using a confocal light microscope with a 100× magnification lens to obtain the grid atlas. The EM grids with cultured cells were then gently picked up with tweezers, and 4 μL of nanogold particle solution (10 nm in diameter; Aurion, the Netherlands) was added onto each grid. Grids were then transferred into a Leica EM GP for cryo-EM specimen preparation. The Leica EM GP was set to 37 °C and 90% humidity, with a blotting time of 8-10 seconds, and grids were blotted from the back side, followed by plunge-freezing in liquid ethane. Finally, the grids were stored in liquid nitrogen, ready for loading into the electron microscope.

For preparation of isolated retraction fibers for cryo-EM analysis, 3.5 μL sample solution was applied onto the glow-discharged Quantifoil grids (Au, R1.2/1.3, 300 mesh). Grids were then loaded into a Vitrobot Mark IV (Thermo Fisher Scientific) with the chamber set to 8°C temperature and 100% humidity. After blotting with filter paper for 4 s at a force of −1, grids were plunge-frozen into liquid ethane and stored in liquid nitrogen for cryo-EM imaging.

### Cryo-electron tomography for the statistical characterization of retraction fiber types

Cryo-ET tilt series of retraction fibers were collected on a Titan Krios microscope operated at 300 kV (Thermo Fisher Scientific) equipped with a K3 direct detector camera (Gatan). Data were recorded in dose-fractionation mode using SerialEM software^7^. Tilt series were collected using a dose-symmetric scheme, with a calibrated pixel size of 1.25 Å, a defocus of −6 μm, a tilt increment of 3°, and an imaging dose of 3 e^−^/Å^2^ for every tilt. The tilt range was from +60° to −60°, starting from 0°, resulting in a total dose of ~123 e^−^/Å^2^. The number of tilt series collected was 12 for the *in situ* TSPAN4-GFP retraction fibers, 76 for the *in situ* TSPAN7-GFP retraction fibers, 75 for the LatA-treated *in situ* TSPAN7-GFP retraction fibers, and 45 for the isolated TSPAN7-GFP retraction fibers. The tilt series were imported to Warp software^8^ for motion correction as well as CTF estimation. The corrected tilt series were firstly aligned in IMOD^9^ using a combination of patch-tracking and gold fiducial markers for tracking. Aligned images were binned by a factor of 8 to a pixel size of 10 Å to generate the tomograms.

### Subtomogram averaging of retraction fibers containing TSPAN7-GFP or engineered TSPAN7 dimers

For subtomogram averaging, 103 tilt series of TSPAN7-GFP-containing retraction fibers and 114 tilt series of engineered TSPAN7 dimer-containing retraction fibers were acquired on a Titan Krios microscope operated at 300 kV (Thermo Fisher Scientific) equipped with a GIF Quantum post-column energy filter (Gatan) and a K3 direct detector camera (Gatan). Tilt series were collected using a dose-symmetric scheme^10^, with a calibrated pixel size of 1.36 Å, a defocus range of −4.5 to −5.5 μm, a tilt increment of 3°, and an imaging dose of 3.2 e^−^/Å^2^ for every tilt. The tilt range was from +60° to −60°, starting from 0°, resulting in a total dose of ~ 131.2 e^−^/Å^2^. Tilt series data preprocessing was conducted in Warp software^8^. Aligned images were binned by a factor of 4 to a pixel size of 5.44 Å, and the flattened tomograms were reconstructed by IMOD^9^.

For particle picking, the retraction fibers were first manually traced in the Dynamo package^11^ using the Dynamo filament model with torsion type. The extreme points of straight segments along each retraction fiber were manually selected to define the central line “backbone”. Bent regions of the retraction fibers were discarded. The parameters *subunits_dz* and *subunits_dphi* were set to 12 pixels and 30°, respectively, to account for the helical rise of the retraction fibers and to minimize the effect of distortions from missing wedges in subtomogram averaging^12^. Sub-volumes with a box size of 128^3^ pixels were cropped using Dynamo with the model-defined positions and initial Euler angles.

Subtomogram averaging and refinement were performed reference-free using Dynamo. A cylindrical mask was generated in Dynamo as an alignment mask, and the other masks were left as default. The initial reference was generated by averaging subtomograms (4 times binned) with 8 rounds of iterative alignment. Subsequently, the initial aligned particle coordinates were imported into RELION3^13^. These 3D particles were extracted and then projected along the Z axis to 2D particles. 2D classification of these particles was performed and the selected particles were grouped into classes with fine structural details. The 3D particles corresponding to the selected 2D particles were utilized for further 3D classification, applying C6 (for TSPAN7-GFP sample) or C7 (for engineered TSPAN7 dimer sample) symmetry and using the density aligned by Dynamo as the initial model.

### Single-particle cryo-EM data collection and analysis

The single-particle cryo-EM datasets for *in situ* TSPAN4-GFP and TSPAN7-GFP retraction fibers were collected using a Thermo Fisher Scientific Titan Krios TEM (300 kV), equipped with a K3 camera (Gatan). The number of micrographs collected was 99 for TSPAN4 and 103 for TSPAN7. Each micrograph was dose-fractionated into 32 frames with a cumulative dose of 50 e^−^/Å^2^ and a pixel size of 1.56 Å. The micrographs were motion-corrected using MotionCor2^14^ and then imported into RELION^15^ for data processing. CTFFIND4^16^ was used to estimate the contrast transfer function (CTF) parameters. For the subsequent two-dimensional classification (Fig. 3b), 1,502 particles were manually picked from the TSPAN4 samples and 1,570 from the TSPAN7 samples, using identical processing parameters.

Single-particle cryo-EM datasets for isolated TSPAN7-GFP retraction fibers were acquired on a Thermo Fisher Scientific Titan Krios TEM (300 kV), equipped with a K3 camera (Gatan) and a GIF Quantum post-column energy filter. A total of 1,823 micrographs were collected, each dose-fractionated into 32 frames with a cumulative dose of 50 e-/Å² and a pixel size of 1.712 Å. Tube particles were manually picked in RELION^17^, and 116,900 particles with an initial helical rise of 88 Å (10% box size, which is larger than the actual rise of 65 Å) were extracted for two-dimensional classification of the retraction fiber. To focus on the classification analysis of the transmembrane region, the coordinates of the above-picked retraction fibers were shifted from their center along the helical axis to their membrane boundaries, and then 361,800 particles were extracted for subsequent 2D classification.

Single-particle cryo-EM datasets of isolated retraction fibers containing engineered TSPAN7 dimers were acquired using a Thermo Fisher Scientific Titan Krios transmission electron microscope (TEM) operating at 300 kV, equipped with a Gatan K3 direct electron detector and a GIF Quantum post-column energy filter. A total of 6,171 micrographs were collected, each dose-fractionated into 32 frames with a cumulative dose of 50 e⁻/Å² and a pixel size of 1.36 Å. Motion correction of the frame stacks was performed using MotionCor2^14^. Dose-weighted micrographs were imported into cryoSPARC^18^ for patch-based contrast transfer function (CTF) estimation. Filament particles were then manually picked in RELION^17^, and 320,972 particles were extracted using an inter-box distance of 66 Å (corresponding to the helical rise). Particle coordinates were then transferred to cryoSPARC for re-extraction and further processing. After multiple rounds of 2D classification, low-quality particles were discarded, yielding 220,083 particles for heterogeneous refinement. Using the sub-tomogram average map as a “good” reference and featureless tubes as a “bad” reference, three rounds of heterogeneous refinement were performed, further eliminating suboptimal particles. The final set of 105,167 particles was subjected to non-uniform (NU) refinement, producing a reconstruction of the spiral structure at 10.0 Å resolution. Precise helical symmetry parameters were then determined using HELIS^19^, yielding a rise of 66.9 Å and a twist of 6.78° for subsequent helical refinement. The helical refinement achieved a 7.8 Å resolution reconstruction of the spiral structure. C7 symmetry expansion was then performed followed by local refinement using a tetramer-focused mask, resulting in a TSPAN7 tetramer structure at 6.6 Å resolution with imposed C2 symmetry. Further C2 symmetry expansion and local refinement with an interdimer-focused mask yielded a dimer map at 5.9 Å resolution.

For model building, the TSPAN7 structure was first predicted using AlphaFold3^20^, then real-space refinement was performed in Phenix^21^ using both the predicted model and our 5.9 Å cryo-EM density to generate the refined TSPAN7 model. This model was then used to build the tetramer and spiral assemblies by docking into the corresponding densities in ChimeraX^22^.

### Statistical analyses and reproducibility

General statistical analyses were performed using GraphPad Prism 8. The statistical results in **Figs. 1b, c, f, j, 2d, 5f, g** and **Extended Data Fig. 1c** were analyzed in and presented by SuperPlots^23^. Data from three independent experiments (colored in yellow, blue and gray) were pooled, and mean (triangles) ± SD were plotted. *P* values of the mean were calculated using a two-tailed paired nonparametric test. In **Figs. 6c, f** and **Extended Data Fig. 9c**, P values were calculated using a two-tailed unpaired nonparametric test. *P* value < 0.05 was considered statistically significant. **** *P* < 0.0001. In all experiments, samples and microscopic images were acquired randomly. The investigators were not blinded to treatment allocation during experiments or to the outcome assessment.

## Acknowledgements

This research was supported by the Ministry of Science and Technology of the People’s Republic of China (grant no. 2024YFA1307301, 2024YFF1502900), the National Natural Science Foundation of China (grant no. 32030023, 92354306, and 32330025 to L.Y., 32130054 to H.-W.W.), Tsinghua-Toyota Joint Research Fund (grant no. 20233930058), Scientific and Technological Innovation Project of China Academy of Chinese Medical Sciences (grant no.CI2023C024YL), Tsinghua University Dushi Program (grant no. 20241080003), the Research Grants Council of Hong Kong (ECS27104125 to N.L.) and The University of Hong Kong’s Seed Fund to N.L.. We thank the State Key Laboratory of Membrane Biology and SLSTU-Nikon Biological Imaging Center for facility support in microscopic imaging and data processing. We thank the Cryo-EM and High-Performance Computation platforms of Tsinghua University Branch of the National Protein Science Facility, and Shuimu BioSciences Ltd. (Beijing) for support in cryo-EM data and SEM data collection. We thank the Laboratory of Animal Resources Center (THU-LARC) of Tsinghua University for technical support. We thank the Cell Biology Facility in Tsinghua University for support in SEM sample preparation. We thank Mr. Xihang Chen (Molecular-Machines&Industries GmbH) for technical support in the optical tweezer system. Sincere gratitude is expressed towards Dr. Sai Li and Mr. Chenhui Zhang for their assistance with cryo-ET data processing.

## Author contributions

L.Y. and H.-W.W. supervised the project and conceived the experiments. D.J.W. conducted cell experiments, animal experiments, imaging, mutation analysis and statistical analysis. D.J.W. and Y.Z. conducted the in vitro reconstitution assays. D.J.W., T.S. and Y.Z. conducted plasmid constructions. X.J., J.R., D.J.W., M.H., X.L., F.Y., N.L. and H-W.W. conducted the cryo-EM data collection and structural determination. X.J., N.L., D.J.W., Q.Z., K.X, and Q.C.Z. performed the model building. R.D. and R. S. designed and performed the micropipette - optical tweezer experiments. S.Y.L. and D.J.W. conducted flow cytometry assays. L.Y., H-W.W., D.J.W., and N.L. wrote the manuscript with input from all authors.

## Data availability

The subtomogram averaging results for retraction fibers with TSPAN7-GFP and the engineered TSPAN7 dimer have been deposited in the Electron Microscopy Data Bank (EMDB) under EMD-65524 and EMD-65527, respectively. Single-particle cryo-EM reconstructions are also available in the EMDB and PDB, including the engineered TSPAN7 dimer spiral (EMD-65483), locally refined TSPAN7 tetramer (EMD-65484; PDB: 9W2D), and locally refined TSPAN7 dimer (EMD-65485; PDB: 9W2B). Additional data and materials can be obtained from the corresponding authors upon request.

## Additional information

Supplementary Information is available for this paper.

Correspondence and requests for materials should be addressed to L.Y.

## Main references

1 Mattila, P. K. & Lappalainen, P. Filopodia: molecular architecture and cellular functions. Nature Reviews Molecular Cell Biology 9, 446–454, doi:10.1038/nrm2406 (2008).

2 Zeng, H. & Sanes, J. R. Neuronal cell-type classification: challenges, opportunities and the path forward. Nature Reviews Neuroscience 18, 530–546, doi:10.1038/nrn.2017.85 (2017).

3 Rustom, A., Saffrich, R., Markovic, I., Walther, P. & Gerdes, H.-H. Nanotubular Highways for Intercellular Organelle Transport. Science 303, 1007–1010, doi:10.1126/science.1093133 (2004).

4 Taylor, A. C. & Robbins, E. Observations on microextensions from the surface of isolated vertebrate cells. Developmental Biology 7, 660–673, 10.1016/0012-1606(63)90150-7 (1963).

5 Buszczak, M., Inaba, M. & Yamashita, Y. M. Signaling by Cellular Protrusions: Keeping the Conversation Private. Trends in Cell Biology 26, 526–534, doi:10.1016/j.tcb.2016.03.003 (2016).

6 Jacquemet, G., Hamidi, H. & Ivaska, J. Filopodia in cell adhesion, 3D migration and cancer cell invasion. Current Opinion in Cell Biology 36, 23–31, 10.1016/j.ceb.2015.06.007 (2015).

7 Lee, S. Y. et al. Retraction fibers produced by fibronectin-integrin α5β1 interaction promote motility of brain tumor cells.

8 Mill, P., Christensen, S. T. & Pedersen, L. B. Primary cilia as dynamic and diverse signalling hubs in development and disease. Nature Reviews Genetics 24, 421–441, doi:10.1038/s41576-023-00587-9 (2023).

9 Kornberg, T. B. & Roy, S. Cytonemes as specialized signaling filopodia. Development 141, 729–736, doi:10.1242/dev.086223 (2014).

10 Pontes, B. et al. Cell Cytoskeleton and Tether Extraction. Biophysical Journal 101, 43–52, doi:10.1016/j.bpj.2011.05.044 (2011).

11 Mahapatra, A. A.-O., Uysalel, C. & Rangamani, P. A.-O. The Mechanics and Thermodynamics of Tubule Formation in Biological Membranes.

12 Legant, W. R. et al. Measurement of mechanical tractions exerted by cells in three-dimensional matrices. Nature Methods 7, 969–971, doi:10.1038/nmeth.1531 (2010).

13 Roux, A. The physics of membrane tubes: soft templates for studying cellular membranes. Soft Matter 9, 6726–6736, doi:10.1039/C3SM50514F (2013).

14 Kiesel, P. et al. The molecular structure of mammalian primary cilia revealed by cryo-electron tomography. Nature Structural & Molecular Biology 27, 1115–1124, doi:10.1038/s41594-020-0507-4 (2020).

15 Conde, C. & Cáceres, A. Microtubule assembly, organization and dynamics in axons and dendrites. Nature Reviews Neuroscience 10, 319–332, doi:10.1038/nrn2631 (2009).

16 Xu, K., Zhong, G. & Zhuang, X. Actin, Spectrin, and Associated Proteins Form a Periodic Cytoskeletal Structure in Axons. Science 339, 452–456, doi:10.1126/science.1232251 (2013).

17 Blumenthal, A. et al. Morphology and migration of podocytes are affected by CD151 levels. American Journal of Physiology-Renal Physiology 302, F1265–F1277, doi:10.1152/ajprenal.00468.2011 (2012).

18 Runge, K. E. et al. Oocyte CD9 is enriched on the microvillar membrane and required for normal microvillar shape and distribution. Developmental Biology 304, 317–325, 10.1016/j.ydbio.2006.12.041 (2007).

19 Bari, R. et al. Tetraspanins regulate the protrusive activities of cell membrane. Biochemical and Biophysical Research Communications 415, 619–626, doi:10.1016/j.bbrc.2011.10.121 (2011).

20 Dharan, R. & Sorkin, R. Tetraspanin proteins in membrane remodeling processes. Journal of Cell Science 137, jcs261532, doi:10.1242/jcs.261532 (2024).

21 Dharan, R. et al. Transmembrane proteins tetraspanin 4 and CD9 sense membrane curvature. Proceedings of the National Academy of Sciences 119, e2208993119, doi:10.1073/pnas.2208993119 (2022).

22 Hemler, M. E. Tetraspanin functions and associated microdomains. Nature Reviews Molecular Cell Biology 6, 801–811, doi:10.1038/nrm1736 (2005).

23 Pöge, M. et al. Determinants shaping the nanoscale architecture of the mouse rod outer segment. eLife 10, e72817, doi:10.7554/eLife.72817 (2021).

24 Oostergetel, G. T., Keegstra, W. & Brisson, A. Structure of the major membrane protein complex from urinary bladder epithelial cells by cryo-electron crystallography1 1Edited by W. Baumeister. Journal of Molecular Biology 314, 245–252, 10.1006/jmbi.2001.5128 (2001).

25 Min, G., Zhou, G., Schapira, M., Sun, T.-T. & Kong, X.-P. Structural basis of urothelial permeability barrier function as revealed by Cryo-EM studies of the 16 nm uroplakin particle. Journal of Cell Science 116, 4087–4094, doi:10.1242/jcs.00811 (2003).

26 Holinski-Feder, E., et al. Nonsyndromic X-linked mental retardation: mapping of MRX58 to the pericentromeric region.

27 Abidi Fe Fau - Holinski-Feder, E. et al. A novel 2 bp deletion in the TM4SF2 gene is associated with MRX58.

28 Zemni, R. et al. A new gene involved in X-linked mental retardation identified by analysis of an X;2 balanced translocation.

29 Ménager, Mickaël M. & Littman, Dan R. Actin Dynamics Regulates Dendritic Cell-Mediated Transfer of HIV-1 to T Cells. Cell 164, 695–709, doi:10.1016/j.cell.2015.12.036 (2016).

30 McLaughlin, K. A., Tombs, M. A. & Christie, M. R. Autoimmunity to tetraspanin-7 in type 1 diabetes. Medical Microbiology and Immunology 209, 437–445, doi:10.1007/s00430-020-00674-2 (2020).

31 Takagi, S. et al. Identification of a highly specific surface marker of T-cell acute lymphoblastic leukemia and neuroblastoma as a new member of the transmembrane 4 superfamily. International Journal of Cancer 61, 706–715, 10.1002/ijc.2910610519 (1995).

32 Cheong, C. M. et al. Tetraspanin 7 (TSPAN7) expression is upregulated in multiple myeloma patients and inhibits myeloma tumour development in vivo. Experimental Cell Research 332, 24–38, 10.1016/j.yexcr.2015.01.006 (2015).

33 Bassani, S. et al. The X-Linked Intellectual Disability Protein TSPAN7 Regulates Excitatory Synapse Development and AMPAR Trafficking. Neuron 73, 1143–1158, 10.1016/j.neuron.2012.01.021 (2012).

34 Jacobson, K., Liu, P. & Lagerholm, B. C. The Lateral Organization and Mobility of Plasma Membrane Components. Cell 177, 806–819, 10.1016/j.cell.2019.04.018 (2019).

35 Tamura, K., Stecher, G. & Kumar, S. A.-O. MEGA11: Molecular Evolutionary Genetics Analysis Version 11.

36 Dharan, R. et al. Intracellular pressure controls the propagation of tension in crumpled cell membranes. Nature Communications 16, 91, doi:10.1038/s41467-024-55398-1 (2025).

37 Abramson, J. et al. Accurate structure prediction of biomolecular interactions with AlphaFold 3. Nature 630, 493–500, doi:10.1038/s41586-024-07487-w (2024).

## Methods references

1 Zheng, L., Baumann, U. & Reymond, J. L. An efficient one-step site-directed and site-saturation mutagenesis protocol. Nucleic Acids Res 32, doi:ARTN e115 10.1093/nar/gnh110 (2004).

2 Nelson, M.D., and Fitch, D.H.A. (2011). Overlap Extension PCR: An Efficient Method for Transgene Construction. In Molecular Methods for Evolutionary Genetics, V. Orgogozo, and M.V. Rockman, eds. (Humana Press), pp. 459–470. 10.1007/978-1-61779-228-1_27.

3 Conchinha, N. V. et al. Protocols for endothelial cell isolation from mouse tissues: brain, choroid, lung, and muscle. STAR Protocols 2, 100508, 10.1016/j.xpro.2021.100508 (2021).

4 Alexandre, Y. O. & Mueller, S. N. Isolation and Analysis of Stromal Cell Populations from Mouse Lymph Nodes. Bio-protocol 7, e2445, doi:10.21769/BioProtoc.2445 (2017).

5 Dharan, R. et al. Transmembrane proteins tetraspanin 4 and CD9 sense membrane curvature. Proceedings of the National Academy of Sciences 119, e2208993119, doi:10.1073/pnas.2208993119 (2022).

6 Zheng, L. et al. Uniform thin ice on ultraflat graphene for high-resolution cryo-EM. Nature Methods 20, 123–130, doi:10.1038/s41592-022-01693-y (2023).

7 Mastronarde, D. N. Automated electron microscope tomography using robust prediction of specimen movements. Journal of Structural Biology 152, 36–51, 10.1016/j.jsb.2005.07.007 (2005).

8 Tegunov, D. & Cramer, P. Real-time cryo-electron microscopy data preprocessing with Warp. Nature Methods 16, 1146–1152, doi:10.1038/s41592-019-0580-y (2019).

9 Kremer, J. R., Mastronarde, D. N. & McIntosh, J. R. Computer Visualization of Three-Dimensional Image Data Using IMOD. Journal of Structural Biology 116, 71–76, 10.1006/jsbi.1996.0013 (1996).

10 Hagen, W. J. H., Wan, W. & Briggs, J. A. G. Implementation of a cryo-electron tomography tilt-scheme optimized for high resolution subtomogram averaging. Journal of Structural Biology 197, 191–198, 10.1016/j.jsb.2016.06.007 (2017).

11 Castaño-Díez, D., Kudryashev, M., Arheit, M. & Stahlberg, H. Dynamo: A flexible, user-friendly development tool for subtomogram averaging of cryo-EM data in high-performance computing environments. Journal of Structural Biology 178, 139–151, 10.1016/j.jsb.2011.12.017 (2012).

12 Zhang, X. et al. Molecular mechanisms of stress-induced reactivation in mumps virus condensates. Cell 186, 1877–1894.e1827, doi:10.1016/j.cell.2023.03.015 (2023).

13 Bharat, Tanmay A. M., Russo, Christopher J., Löwe, J., Passmore, Lori A. & Scheres, Sjors H. W. Advances in Single-Particle Electron Cryomicroscopy Structure Determination applied to Sub-tomogram Averaging. Structure 23, 1743–1753, 10.1016/j.str.2015.06.026 (2015).

14 Zheng, S. Q. et al. MotionCor2: anisotropic correction of beam-induced motion for improved cryo-electron microscopy. Nature Methods 14, 331–332, doi:10.1038/nmeth.4193 (2017).

15 Zivanov, J. A.-O. et al. New tools for automated high-resolution cryo-EM structure determination in RELION-3. LID - 10.7554/eLife.42166 [doi] LID - e42166.

16 Rohou, A. & Grigorieff, N. CTFFIND4: Fast and accurate defocus estimation from electron micrographs. Journal of Structural Biology 192, 216–221, doi:10.1016/j.jsb.2015.08.008 (2015).

17 He, S. & Scheres, S. H. W. Helical reconstruction in RELION. Journal of Structural Biology 198, 163–176, 10.1016/j.jsb.2017.02.003 (2017).

18 Punjani, A., Rubinstein, J. L., Fleet, D. J. & Brubaker, M. A. cryoSPARC: algorithms for rapid unsupervised cryo-EM structure determination. Nature Methods 14, 290–296, doi:10.1038/nmeth.4169 (2017).

19 Huang, M. et al. Accurate helical parameter estimation based on cylindrical unrolling. Structure 33, 1–11, 10.1016/j.str.2025.06.008 (2025).

20 Abramson, J. et al. Accurate structure prediction of biomolecular interactions with AlphaFold 3. Nature 630, 493–500, doi:10.1038/s41586-024-07487-w (2024).

21 Liebschner, D. et al. Macromolecular structure determination using X-rays, neutrons and electrons: recent developments in Phenix. Acta Crystallographica Section D 75, 861–877, doi:doi:10.1107/S2059798319011471 (2019).

22 Meng, E. C. et al. UCSF ChimeraX: Tools for structure building and analysis. Protein Science 32, e4792, 10.1002/pro.4792 (2023).

23 Lord, S. J., Velle, K. B., Mullins, R. D. & Fritz-Laylin, L. K. SuperPlots: Communicating reproducibility and variability in cell biology. Journal of Cell Biology 219, e202001064, doi:10.1083/jcb.202001064 (2020).

