## Extended data for "Polymerization of Tetraspanin 7 into Helical Transmembrane Skeletons for Tubular Membrane Stabilization"

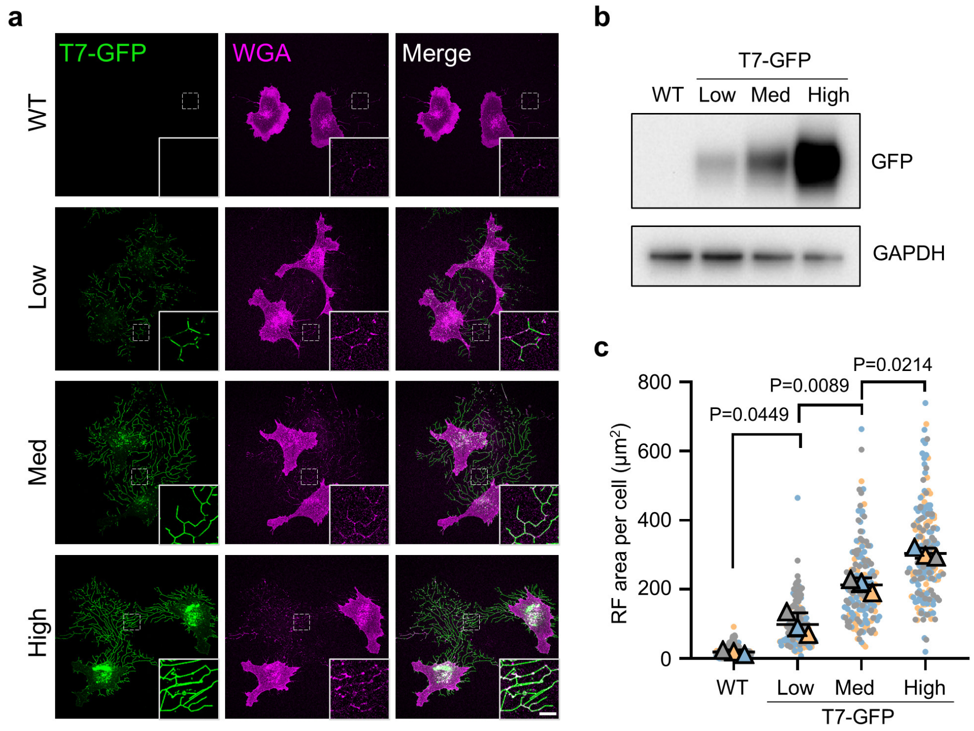


**Extended Data Fig. 1 TSPAN7 promotes the formation of retraction fibers in a dose-dependent manner.**

**a,** Representative confocal images of wild-type NRK cells or NRK cells stably expressing different levels of TSPAN7-GFP (Low, Med, and High). Cells were stained with WGA-TMR.
**b,** Western-blot analysis showing the expression levels of TSPAN7-GFP in the 3 different cell lines from **a**.
**c,** Statistical analysis of the area of retraction fibers per cell in **a**. N = 169, 165, 168 and 180 cells were analyzed for WT, T7-Low, T7-Med, and T7-High, respectively.
Scale bars, 5 μm.

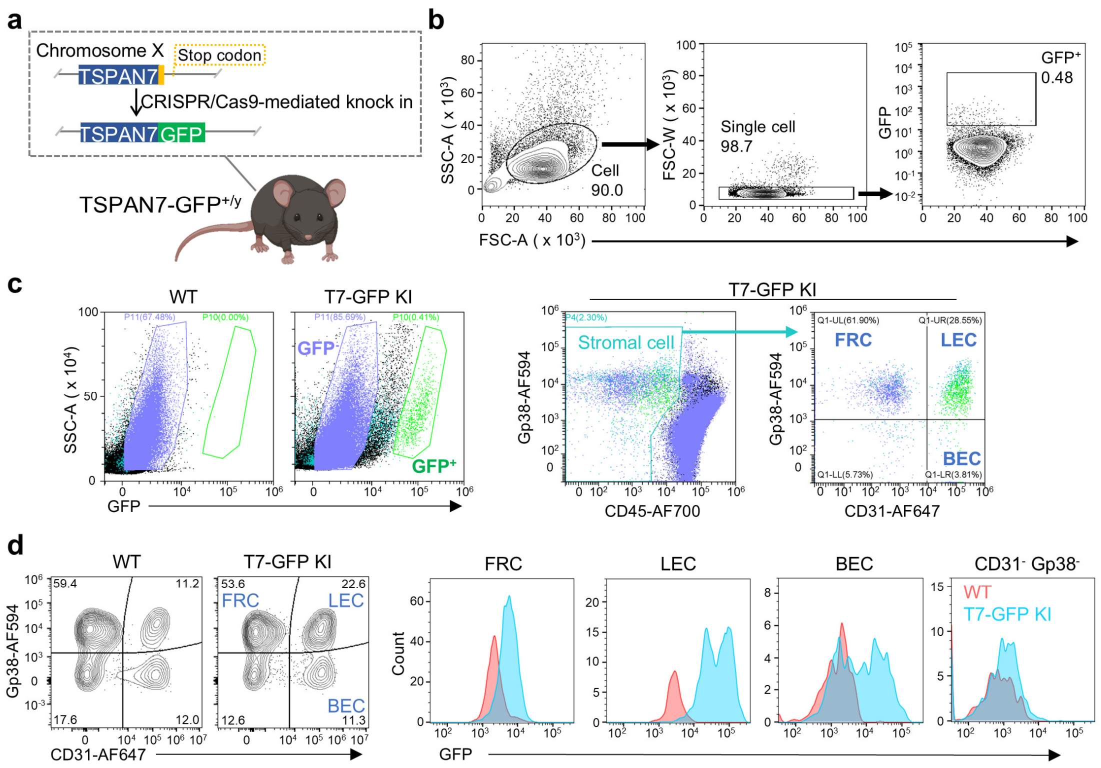


**Extended Data Fig. 2 Identification of GFP⁺ cells in mouse lymph nodes.**

**a,** Schematic illustration of the construction of the TSPAN7-GFP knock-in (T7-GFP KI) mouse strain. The stop codon of TSPAN7 was replaced with the coding sequence of GFP using the CRISPR/Cas9 system.

**b,** The flow cytometry gating strategy that was used in isolating GFP⁺ cells from mouse lymph nodes.

**c,** Representative flow cytometry plots of GFP⁺ cells in lymph nodes from WT and TSPAN7-GFP knock-in mice (left) and the gating strategy for analyzing GFP⁺ lymph node stromal cell populations (right) including FRCs (fibroblastic reticular cells), LECs (lymphatic endothelial cells), and BECs (blood endothelial cells).

**d,** Representative flow cytometry plots (left panel) of lymph node stromal cell populations and mean fluorescence intensity (MFI) of GFP (right panel) in different populations from WT and T7-GFP KI mice.

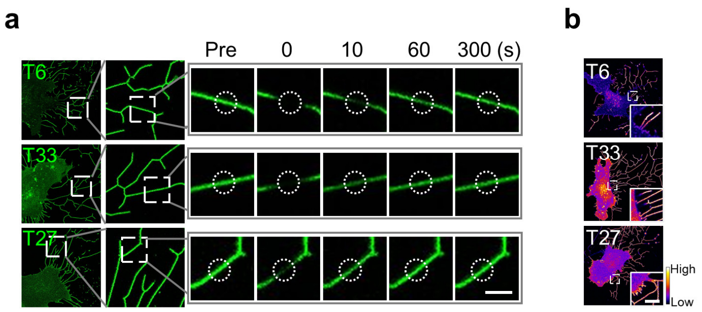


**Extended Data Fig. 3 Analysis of TSPAN6, TSPAN27 and TSPAN33 on retraction fibers, related to Fig. 2.**

**a,** Related to **Fig. 2i**. Representative time-lapse images of FRAP assays on the retraction fibers of NRK cells expressing TSPAN6-GFP (T6), TSPAN27-GFP (T27) or TSPAN33-GFP (T33). The right panels show enlarged areas from the left panels. The dashed circles represent areas of bleaching.
**b,** Related to **Fig. 2j**. The fluorescence intensities of TSPAN6-GFP (T6), TSPAN27-GFP (T27) and TSPAN33-GFP (T33) are shown as heatmaps.
Scale bars, 2 µm in **a**; 5 µm in **b**.

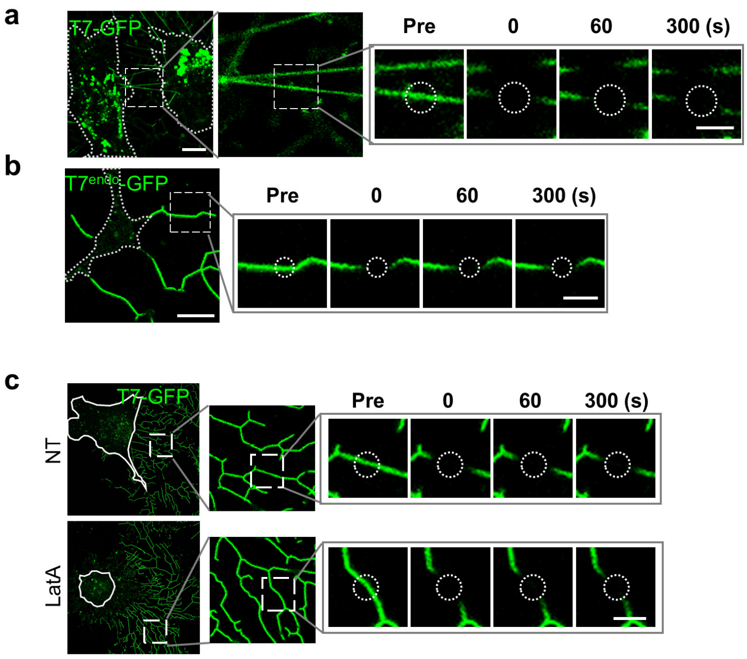


**Extended Data Fig. 4 TSPAN7 is immobile on TNTs and the retraction fibers in LECs, related to Fig. 2.**

**a,** Related to **Fig. 2k**. Representative time-lapse images of FRAP assays on the TNTs of T7-GFP-expressing NRK cells. Cell contours are highlighted with dashed white lines. The right panels show enlarged areas from the left panels. The dashed circles represent areas of bleaching.

**b,** Related to **Fig. 2l**. Representative time-lapse images of FRAP assays on the retraction fibers of $GFP^{+}$ cells isolated from the lymph nodes of TSPAN7-GFP knock-in mice. Cell contours are highlighted with dashed white lines. The right panels show enlarged areas from the left panels. The dashed circles represent areas of bleaching.

**c,** Related to **Fig. 2m**. Confocal images of cells pretreated with 2 μM latrunculin A (LatA) or control solvent (ethanol, NT) before FRAP assay are shown in the left panel. Cell contours are highlighted with white lines. The right panels show enlarged areas from the left panels. The dashed circles represent areas of bleaching.

Scale bars, 10 μm in **a** (left panel); 5 μm in **b** (left panel); 2 μm in enlarged areas in **a**, **b** and **c**.

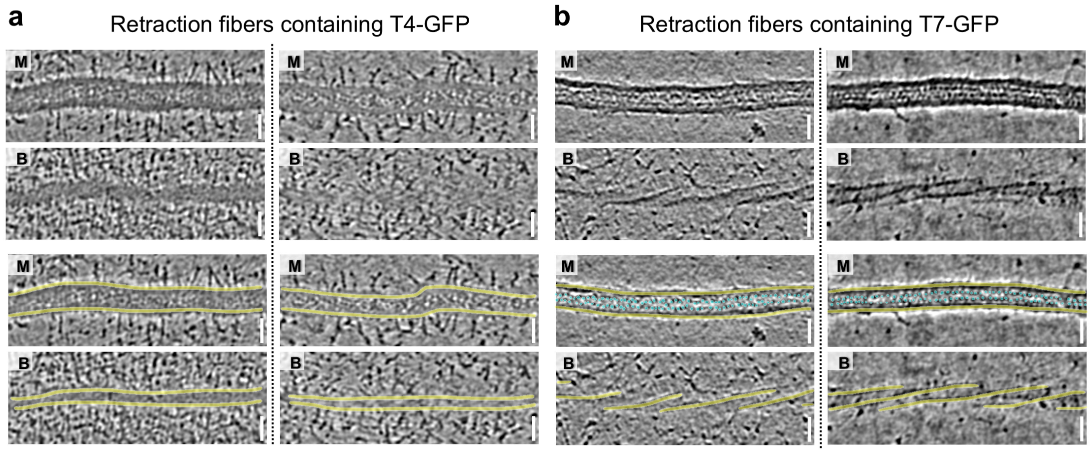


**Extended Data Fig. 5 Representative cryo-ET reconstructions of TSPAN4-GFP and TSPAN7-GFP retraction fibers.**

**a,** The middle (M) and bottom (B) slices of two additional TSPAN4-GFP retraction fiber tomograms are shown. The retraction fiber boundaries in different slices are labeled with yellow lines.
**b,** The middle (M) and bottom (B) slices of two additional TSPAN7-GFP retraction fiber tomograms are shown. The yellow lines outline the retraction fiber boundaries in different slices, and the cyan markers indicate the density of the regular dots (GFP) on the cytoplasmic side of the cell membrane. Scale bars, 30 nm.

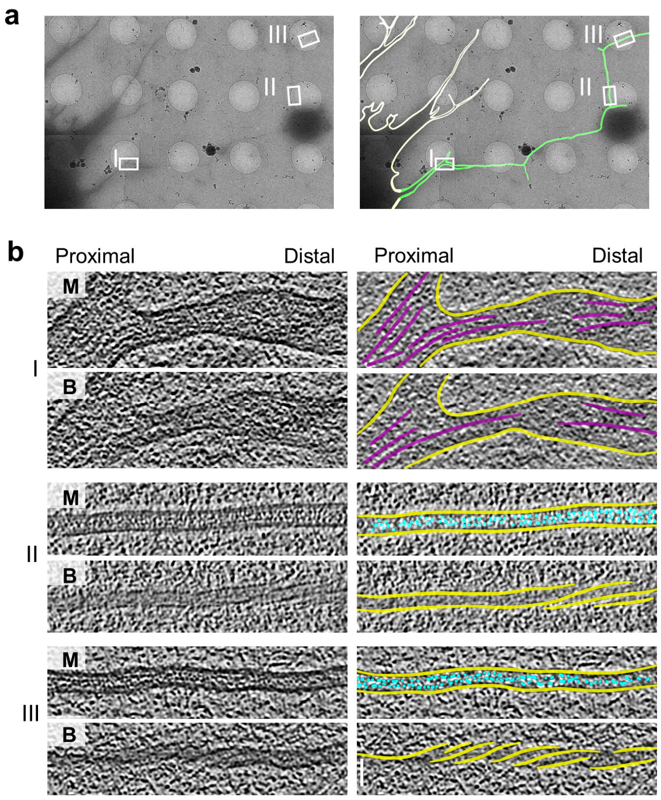


**Extended Data Fig. 6 Cryo-ET characterization of the spiral assembly of TSPAN7 on retraction fibers attached to cells.**

**a,** Left: Representative low-magnification cryo-EM micrograph showing retraction fibers extending from the cell body. Right: The same image with the cell boundary indicated by white lines and retraction fibers indicated by green lines.

**b,** Left: Middle (M) and bottom (B) slices of the tomograms corresponding to the three areas labeled as I-III in **a**. Right: The same images with F-actin labeled in magenta, the cell membrane in yellow, and GFP in cyan.

Scale bar, 30 nm.

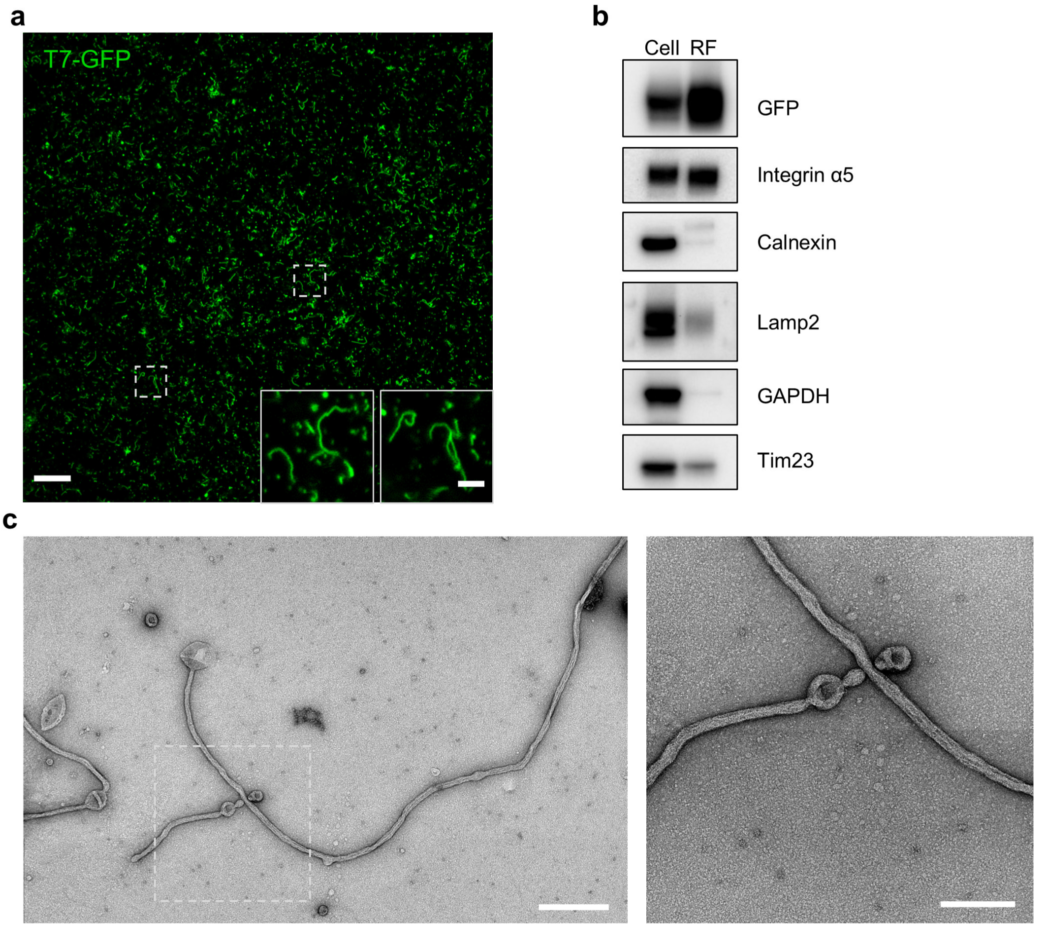


**Extended Data Fig. 7 Characterization of isolated retraction fibers.**

**a,** Representative confocal image of retraction fibers isolated from NRK cells stably expressing TSPAN7-GFP.

**b,** Western-blot analysis of cell bodies (Cell) or retraction fibers (RF). Equal amounts of protein were loaded for the two samples.

**c,** Representative negative-stain EM images of isolated retraction fibers from NRK cells stably expressing TSPAN7-GFP. The right panel shows a magnified view of the region indicated by the white dashed box in the left panel.

Scale bars, 10 μm in **a**; 2 μm in inserts in **a**; 500 nm in the left panel in **c**; 200 nm in the right panel in **c**.

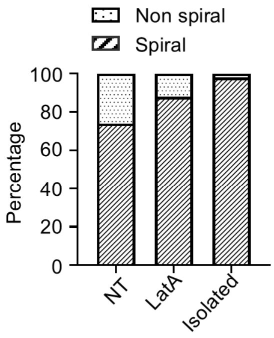


**Extended Data Fig. 8 Prevalence of TSPAN7 spirals on retraction fibers.**

Cryo-EM tilt series were collected from 76 retraction fibers (RFs) from untreated cells (NT), 75 RFs from LatA-treated cells, and 45 isolated RFs. Following tomogram reconstruction, each fiber was classified as either “Non-spiral” or “Spiral”. Quantitative analysis revealed spiral configurations in 74% (NT), 88% (LatA-treated), and 98% (isolated) of cases.

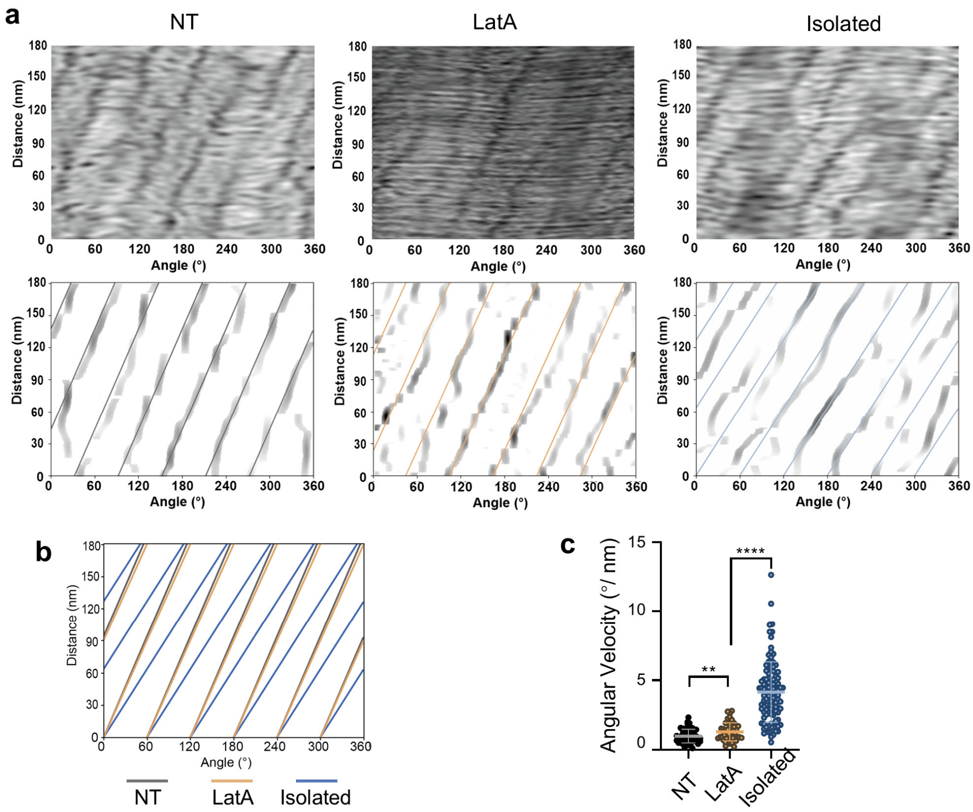


**Extended Data Fig. 9 Statistics of TSPAN7 spiral parameters on retraction fibers.**

**a,** Helical unfolding analysis of representative tomograms of retraction fibers (RFs) from untreated cells (NT), RFs from LatA-treated cells, and isolated RFs. Top: Original unfolded tomograms. Bottom: Corresponding filtered and feature-extracted images, as well as the fitted helical crest.

**b,** Superimposed spiral crest fittings from panel **a**.

**c,** Plotting of the angular velocity of the TSPAN7 spirals to assess their tightness. N=52 randomly selected spiral structures from each of the three groups were analyzed.

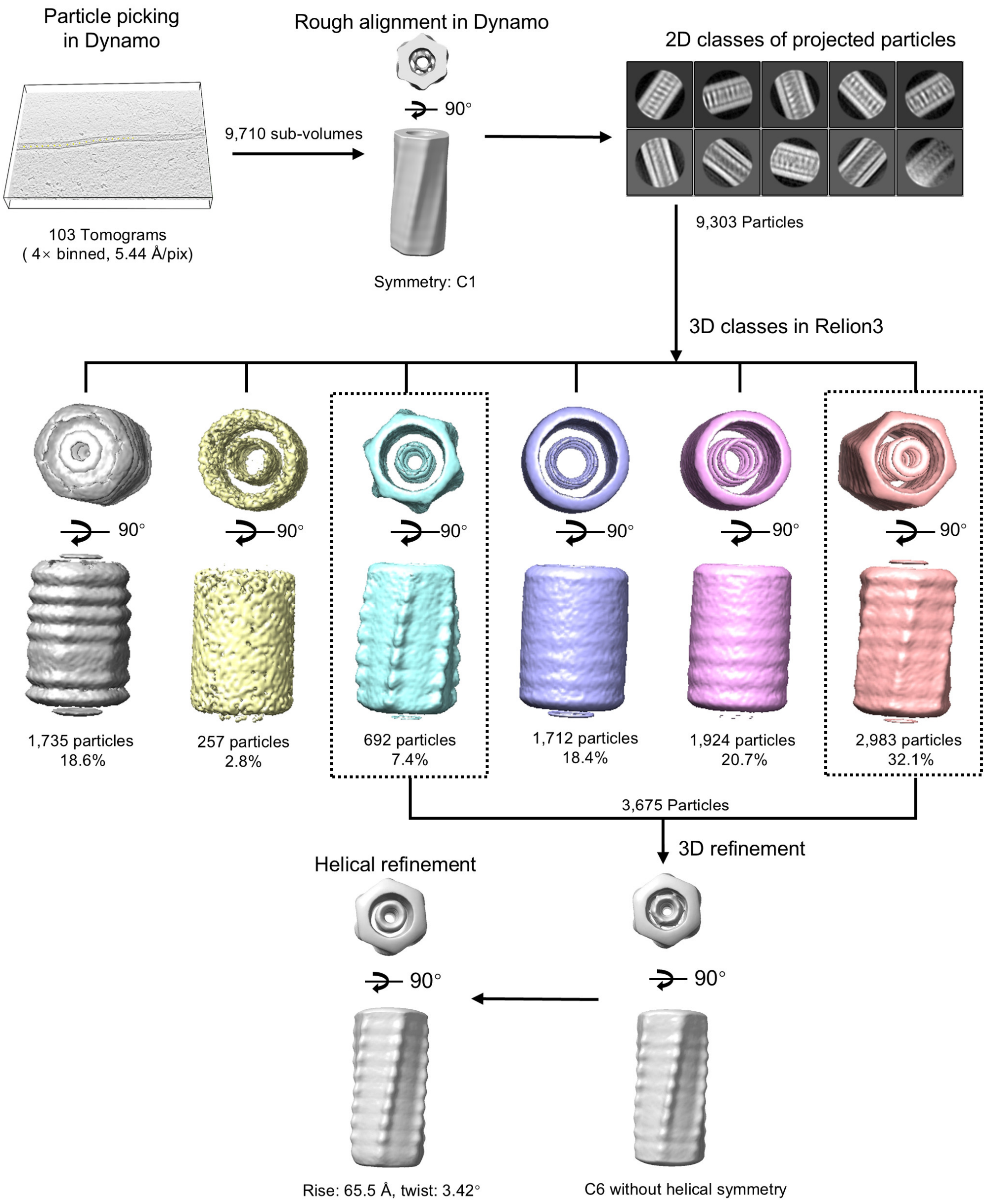


**Extended Data Fig. 10 Workflow of subtomogram averaging analysis for isolated TSPAN7-GFP retraction fibers.**

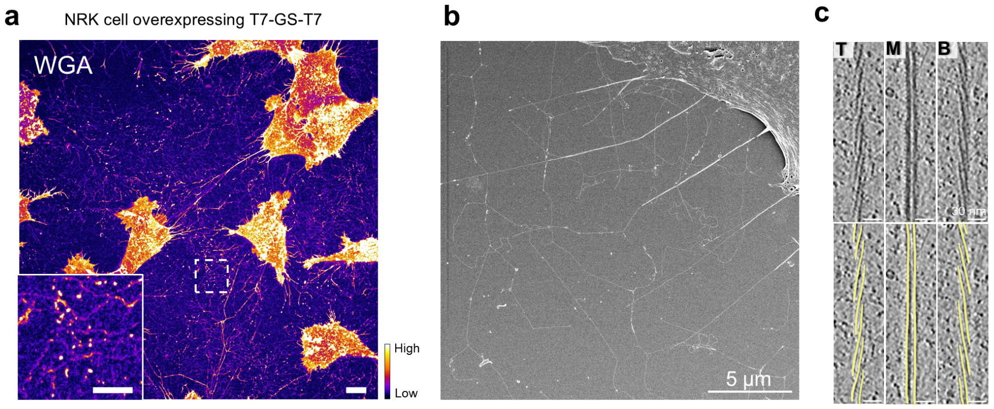


**Extended Data Fig. 11 Characterization of *in situ* retraction fibers containing the engineered TSPAN7 dimer.**

**a,** Representative confocal image of NRK cells stably expressing the engineered TSPAN7 dimer and stained with WGA-TMR. The intensity of the fluorescent signal is shown as a heatmap. Scale bars, 10 μm.

**b,** Representative SEM image of an NRK cell stably expressing the engineered TSPAN7 dimer.

**c,** Representative tomogram slices of the *in situ* retraction fibers containing engineered TSPAN7 dimers. See also **Supplementary video 9**.

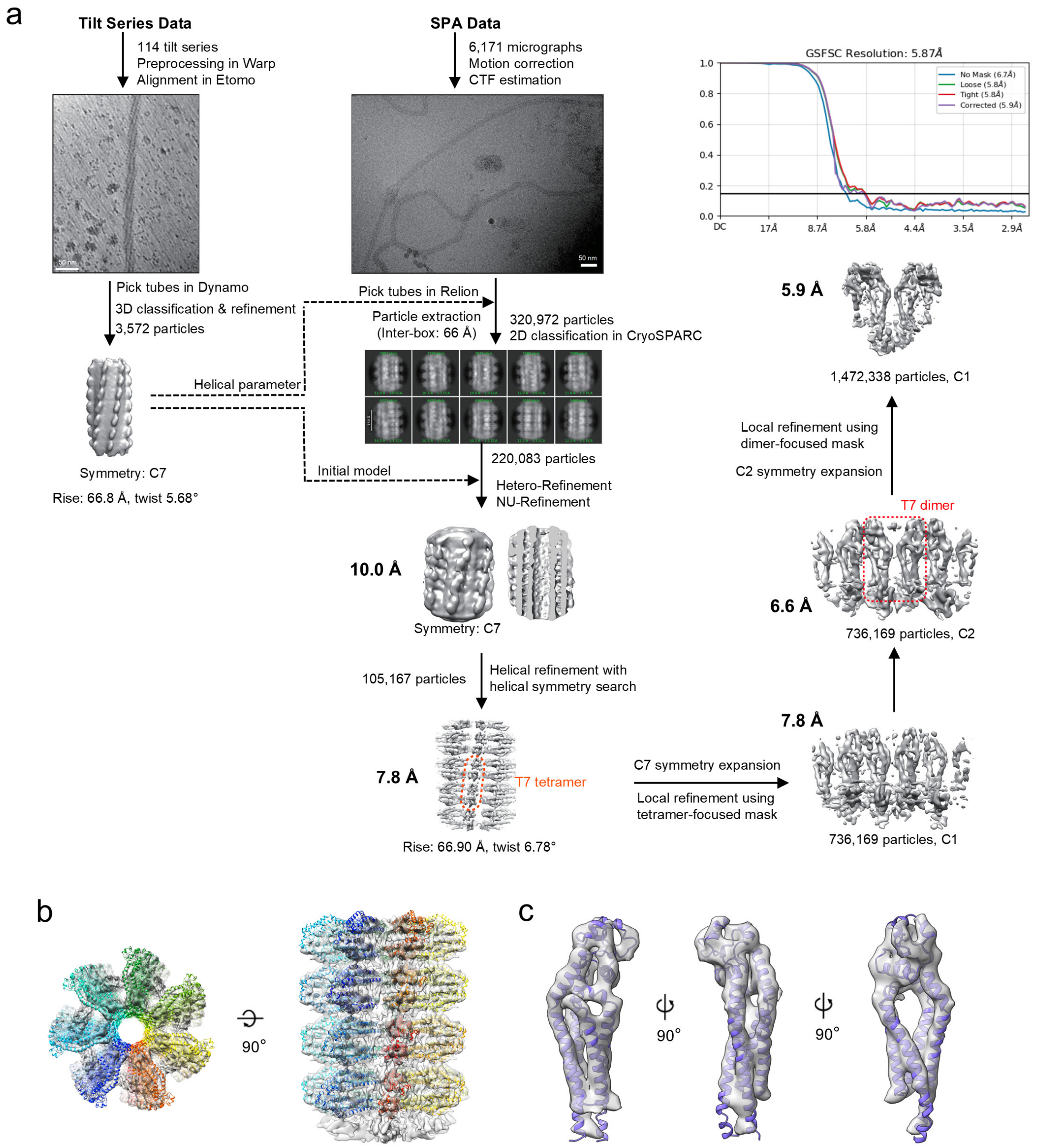


**Extended Data Fig. 12 Cryo-EM reconstruction of retraction fibers containing engineered TSPAN7 dimers.**

**a,** Workflow of cryo-EM data processing.

**b,** TSPAN7 spiral density map with fitted atomic model.

**c,** TSPAN7 monomer density extracted from the spiral map with fitted atomic model.

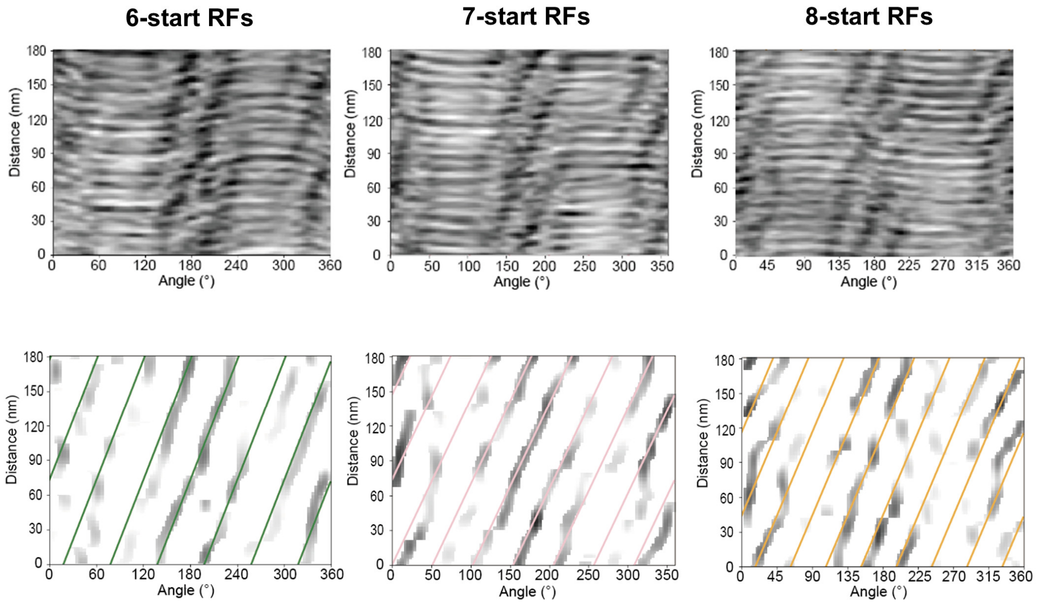


**Extended Data Fig. 13 Helical unfolding analysis of representative tomograms from retraction fibers (RFs) containing engineered TSPAN7 dimers.**

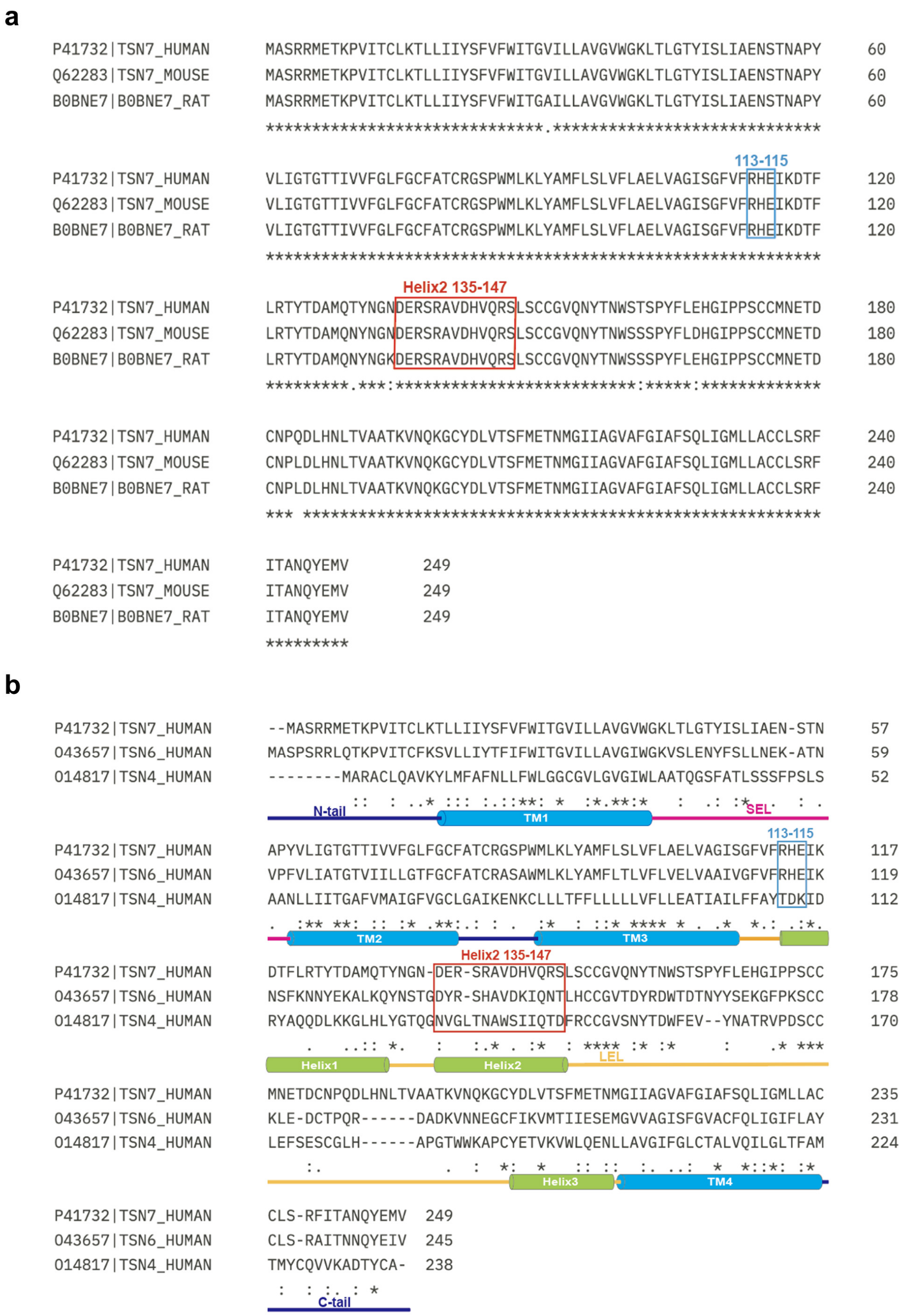


**Extended Data Fig. 14 Tetraspanin sequence alignment.**

**a,** Sequence alignment of TSPAN7 from *H. sapiens*, *M. musculus*, and *R. norvegicus*, conducted with Cluster Omega. Colored boxes mark the two oligomer interaction sites, E115 and Helix 2 (aa 135-147).

**b,** Sequence alignment of human TSPAN7, TSPAN6, and TSPAN4 proteins, conducted with Cluster Omega. Topological domains are indicated below the sequences. Colored boxes mark the two TSPAN7 oligomer interaction sites, E115 and Helix 2 (aa 135-147).

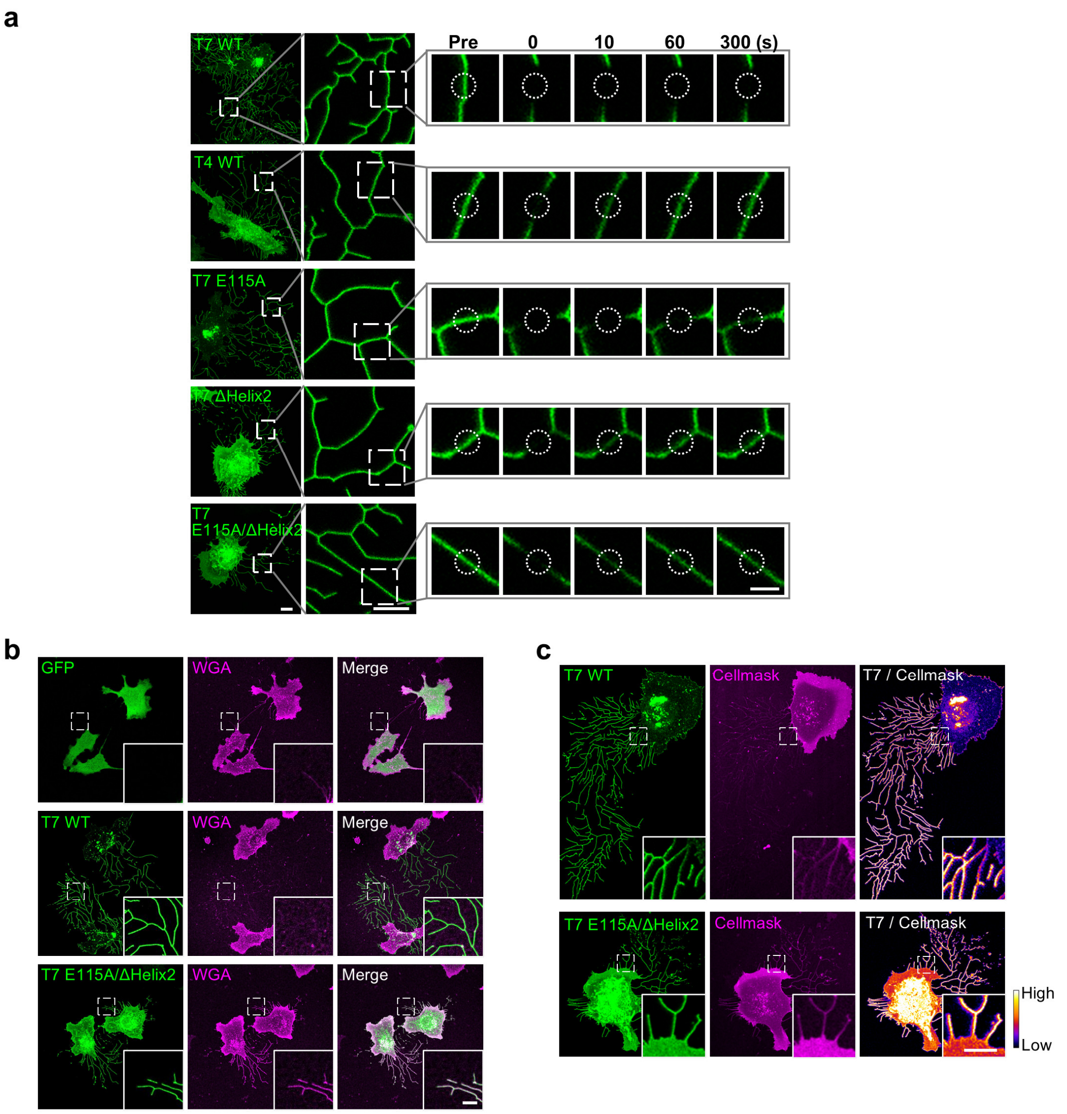


**Extended Data Fig. 15 Analysis of TSPAN7 mutants.**

**a,** Related to **Fig. 5e**. Representative time-lapse images of FRAP assays performed on the retraction fibers of NRK cells expressing wild-type TSPAN7-GFP (T7 WT), wild-type TSPAN4-GFP (T4 WT), single-site mutants of TSPAN7-GFP (T7 E115A or T7 ΔHelix2) and the double mutant of TSPAN7-GFP (T7 E115A/ΔHelix2). The right panels show enlarged areas from the left panels. The dashed circles represent areas of bleaching.

**b,** Related to **Fig. 5f**. Representative confocal images of NRK cells expressing GFP, wild-type TSPAN7-GFP (T7 WT) or double mutant TSPAN7-GFP (T7 E115A/ΔHelix2) and stained with WGA-TMR (WGA).

**c,** Related to **Fig. 5g**. Representative confocal images of NRK cells expressing wild-type TSPAN7-GFP (T7 WT) or double mutant TSPAN7-GFP (T7 E115A/ΔHelix2). The fluorescence intensities of T7 WT and T7 E115A/ΔHelix2 are shown as heatmaps.

Scale bars, 5 μm in **a** (left), **b** and **c**; 2 μm in **a** (right).
