## Supplementary table for "Polymerization of Tetraspanin 7 into Helical Transmembrane Skeletons for Tubular Membrane Stabilization"

**Supplementary Table 1 Cryo-ET data collection and reconstruction statistics**

| | **Retraction fiber with TSPAN7-GFP** | **Retraction fiber with engineered TSPAN7 dimer** |
|---|---|---|
| Magnification | 64,000 | 64,000 |
| Voltage (kV) | 300 | 300 |
| Pixel size (Å) | 1.36 | 1.36 |
| Camera | Gatan K3 Summit | Gatan K3 Summit |
| Tilt range (°) | +60 to -60 | +60 to -60 |
| Tilt step (°) | 3 | 3 |
| Electron exposure ($e^{-}$/Å$^{2}$) | 3.2 (per tilt) | 3.2 (per tilt) |
| Defocus range (μm) | -4.5 to -5.5 | -4.5 to -5.5 |
| Raw tomograms | 103 | 114 |
| Subtomograms | 3,675 | 3,572 |
| Reported resolution (Å) | 33 | 16 |
| Symmetry imposed | C6 | C7 |
| EMDB code | EMD-65524 | EMD-65527 |

**Supplementary Table 2 Single-particle cryo-EM data collection and reconstruction statistics**

| | Retraction fiber with engineered TSPAN7 dimer |
|---|---|
| **Data collection** | |
| Magnification | 64,000 |
| Voltage (kV) | 300 |
| Pixel size (Å) | 1.36 |
| Camera | Gatan K3 Summit |
| Electron exposure ($e^{-}$/$Å^{2}$) | 50 |
| Defocus Range (μm) | -1.2 to -1.8 |
| Number of micrographs | 6,171 |

| **Reconstruction** | **Spiral** (EMD-65483) | **Tetramer** (EMD-65484; PDB: 9W2D) | **Dimer** (EMD-65485 PDB: 9W2B ) |
|---|---|---|---|
| Particles | 105,167 particles | 736,169 particles | 1,472,338 particles |
| Software | CryoSPARC | CryoSPARC | CryoSPARC |
| Reported resolution (Å) (FSC=0.143 threshold) | 7.8 | 6.6 | 5.9 |
| Symmetry imposed | Helical | C2 | C2 |
| **Model statistics** | | | |
| Chains | / | 4 | 2 |
| Atoms | / | 7732 | 3866 |
| Protein residues | / | 1000 | 500 |
| r.m.s. deviations | | | |
| Bonds (Å) | / | 0.002 | 0.002 |
| Angles ($^{o}$) | / | 0.740 | 0.621 |
| Ramachandran plot | | | |
| Favored (%) | / | 96.77 | 97.58 |
| Allowed (%) | / | 3.23 | 2.42 |
| Outliers (%) | / | 0.00 | 0.00 |
| MolProbity score | / | 1.85 | 1.66 |
| Clash score | / | 13.91 | 11.39 |
| Rotamer outliers (%) | / | 0.00 | 0.00 |
